# dATP Elevation Induces Myocardial Metabolic Remodeling to Support Improved Cardiac Function

**DOI:** 10.1101/2022.11.07.515235

**Authors:** Ketaki N Mhatre, Jason D Murray, Galina Flint, Timothy S. McMillen, Gerhard Weber, Majid Shakeri, An-Yue Tu, Sonette Steczina, Robert Weiss, David J. Marcinek, Charles E Murry, Daniel Raftery, Rong Tian, Farid Moussavi-Harami, Michael Regnier

## Abstract

Hallmark features of systolic heart failure are reduced contractility and impaired metabolic flexibility of the myocardium. Cardiomyocytes (CMs) with elevated deoxy ATP (dATP) via overexpression of ribonucleotide reductase (RNR) enzyme robustly improve contractility. However, the effect of dATP elevation on cardiac metabolism is unknown. Here, we developed proteolysis-resistant versions of RNR and demonstrate that elevation of dATP/ATP to ~1% in CMs in a transgenic mouse (TgRRB) resulted in robust improvement of cardiac function. Pharmacological approaches showed that CMs with elevated dATP have greater basal respiratory rates by shifting myosin states to more active forms, independent of its isoform, in relaxed CMs. Targeted metabolomic profiling revealed a significant reprogramming towards oxidative phosphorylation in TgRRB-CMs. Higher cristae density and activity in the mitochondria of TgRRB-CMs improved respiratory capacity. Our results revealed a critical property of dATP to modulate myosin states to enhance contractility and induce metabolic flexibility to support improved function in CMs.

**Highlights:**

- Ubiquitylation-resistant variant RRB in a transgenic mice model (TgRRB) elevates dATP level up to 1% (of the total ATP pool) in the heart and improves function.
- TgRRB-CMs show greater basal oxygen consumption due to changes in myosin state by dATP.
- TgRRB-CMs respond to elevated function with a metabolic shift, such that there are higher pools of oxidative metabolites, with elevated OXPHOS, FAO, and energy reserve.
- Long-term mitochondrial remodeling may occur to accommodate for the higher energy demands of the high functioning TgRRB-CMs.

**Graphical abstract:** 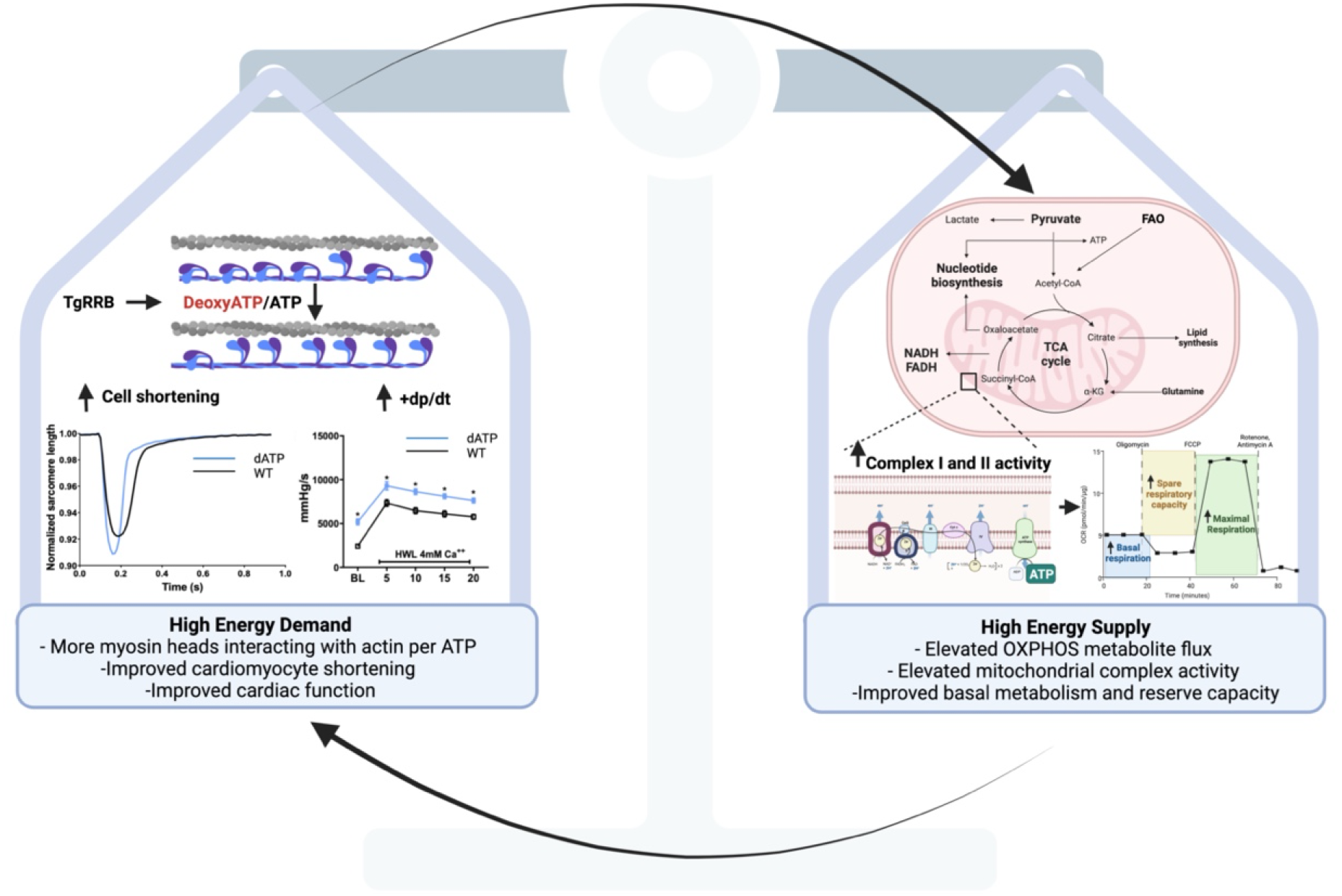

## Introduction

Heart failure (HF) is a clinical diagnosis based on a constellation of symptoms due to impaired ventricular filling or ejection of blood, the latter of which is dependent on actin-myosin cross bridge cycling during systolic muscle contraction. There are also dramatic changes in cardiac metabolism in the failing hearts, such as loss of metabolic flexibility and energy deficiency[1]. Traditional inotropic agents exert their effect by modulating calcium signaling pathways, but their use is associated with poor patient outcomes[2]. All the currently approved positive inotropic drugs increase intracellular cyclic adenosine monophosphate (cAMP) along with cardiac output at the expense of energetic efficiency, which can lead to maladaptive remodeling and arrhythmia[3]. Hence, there is a need for targeted therapies to reduce morbidity and mortality[4–7]. Direct targeting of the myofilament is desirable but many compounds that increase contraction may impair relaxation and cardiac enegetics[8,9]. An ideal therapeutic agent for the treatment of systolic dysfunction would be to improve systolic function and cardiac metabolism without impairing diastolic function.

We have previously reported that the naturally occurring nucleotide dATP is an effective molecule that improves cardiac contractility. dATP binds to cardiac myosin with a similar affinity as ATP, and it enhances the rate and magnitude of both force development and relaxation of cardiac muscle from rodents, porcine, canine, and human tissues[10–15]. This occurs by increasing myosin recruitment[16] and the dynamics of myosin-actin cross bridge cycling[17,18]. Our recent studies showed that enhanced myosin-binding with dATP (vs ATP) occurs via increasing electrostatic interactions between myosin and actin[19,20], as well as by augmented recruitment of myosin from a low activity state (SRX) on the thick filament backbone to a higher activity state (DRX) where one or both heads are free to interact with thin filament[21]. In skeletal muscle, we provided evidence that elevated dATP maintains myosin S1 heads in a position closer to thin filaments during relaxation and that this can facilitate subsequent contraction cycles[22]. Thus, dATP enhancement of cardiac muscle function occurs via multiple mechanisms.

RNR is the enzyme that converts nucleotides to deoxynucleotides, including dATP. Overexpression of RNR using either a transgenic mouse model (TgRR) or viral vectors, have shown that dATP levels can be increased from the baseline of <0.1% to ~ 1% of the ATP pool in cardiomyocytes (CMs). This, in turn, increases CM contraction, left ventricular (LV) developed pressure and both the rate of pressure development (+dP/dt) and decline (-dP/dt) to improve heart function in both small and large animal models of HF[23–25]. RNR is a hetero-tetramer consisting of two subunits: a large subunit of RRM1 dimer which contains the catalytic site, and one of two isoforms of a small subunit, RRM2 or RRM2B, which associate together to form RNR holoenzyme[26,27]. Over-expression of RRM2 was highly effective in improving cardiac function, via elevation of dATP, in our earlier transgenic model (TgRR). However, this subunit is tightly regulated by the ubiquitin-proteasome system across the cell cycle and thus may pose challenges for its stable expression in post-mitotic cells like CMs[28–30]. In this study, we investigated proteolysis-resistant isoforms of RRM2, RRM2B, [28,30] to stably elevate dATP levels in CMs and improve its function.

Recent studies have shown that modulation of the myosin state (based on ATPase turnover) via disease-related mutations[31] not only alters contractile function at the cellular and organ level but also cellular energetics[32]. These mutations can augment recruitment of myosin from a low activity state (SRX) on the thick filament backbone to a higher activity state (DRX) where one or both heads are free to interact with thin filament[21] and can contribute to the observed increase in the energy cost of force generation[32,33]. This can result in energy deficient CMs leading to enhanced ROS production, mitochondria damage, and altered mitochondrial content, cristae, and respiratory complexes. This, in the long-term, causes myocardial energy deprivation in cardiomyopathies[34]. Thus, myosin states and their effect on contractile activity can modulate cardiac metabolism and mitochondria function and it would be important to study the long-term effect of myotropes[35], drugs that directly modulate the contractile machinery, like dATP in this respect.

In the current study, we demonstrate that elevating dATP by overexpression of RRM1 and RRM2B elevates LV systolic and diastolic function. dATP elevation has shown to improve function via increase of cross bridge recruitment and binding to actin (crossbridge formation) and crossbridge cycling[16–18,20]. This results in higher basal ATPase turnover, leading to higher energy demands. Current study suggest that cellular metabolism adapts to greater oxidative capacity and mitochondrial remodeling thereby possibly maintaining (d)ATP supply and energetic reserves in high functioning CMs with chronically elevated dATP.

## Results

### RNRB overexpressing mice have elevated dATP and improved systolic and diastolic function

Here we studied a transgenic mouse line (TgRRB) that globally overexpresses two subunits of ribonucleotide reductase: the large subunit RRM1 and the p53-inducible isoform of the small subunit, RRM2B. Wild-type littermates were used as controls.

We used high-performance liquid chromatography-mass spectrometry (HPLC-MS/MS) to measure total dATP (along with other dNTPs, Fig. S2a) and ATP content. The ratio of dATP to ATP was significantly elevated for hearts of 4-month-old RNRB versus WT littermate mice in the myocardium (Fig. 1a; RNRB = 0.91±0.07% vs WT = 0.01±0.002%), in isolated CMs (Fig. S2b) and in isolated mitochondria (Fig. S2c).

**Fig. 1.**
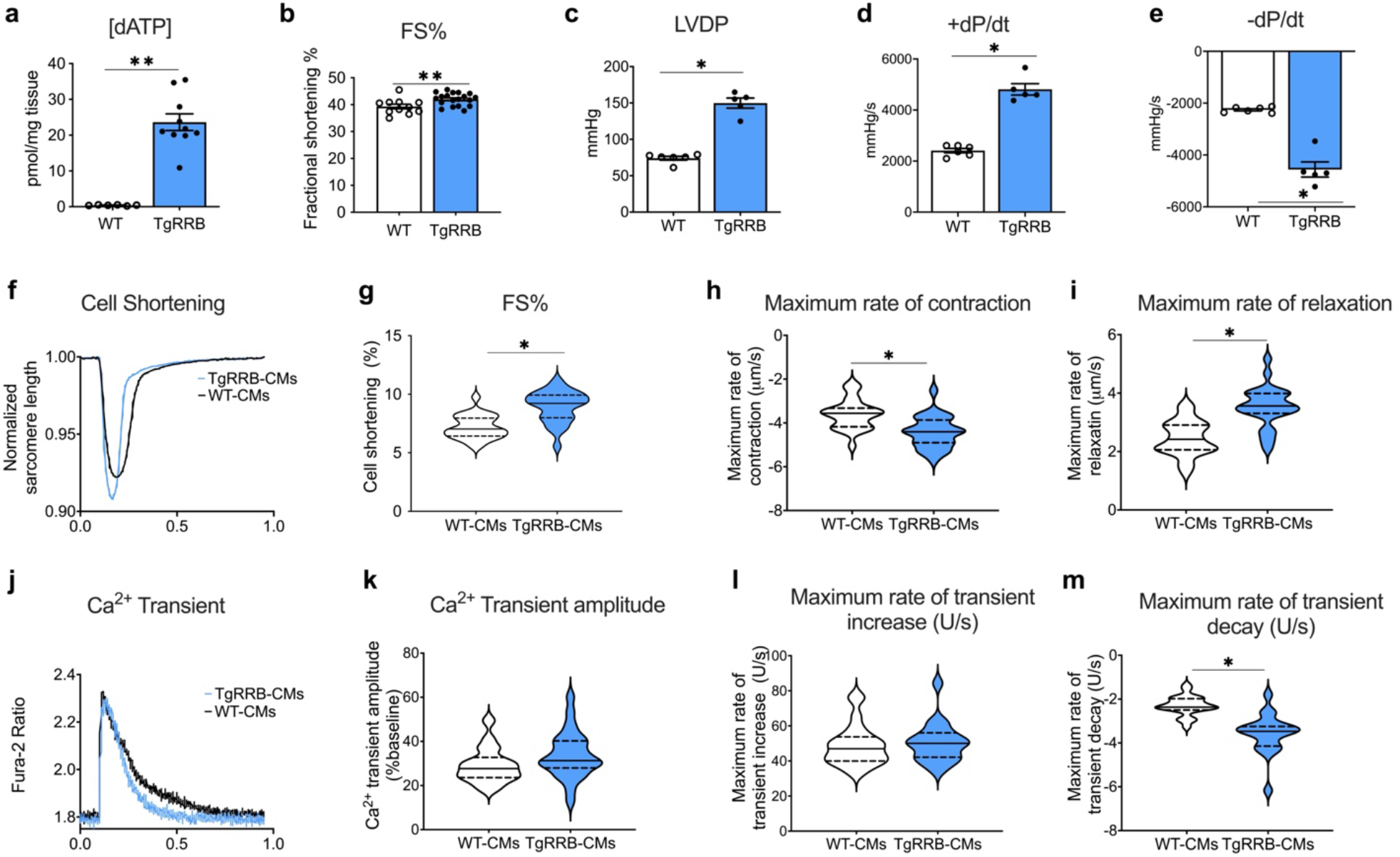
TgRRB mice have elevated dATP and improved systolic and diastolic function. (a) [dATP] mass spectrometry measurements from the myocardium. (b) Echocardiogram measurements FS: Fractional Shortening. (c-e) Baseline contractility in perfused isolated hearts LVDP: Left-ventricular developed pressure, +dP/dt: maximum rate of pressure development in contraction, -dP/dt: maximum rate of pressure development in relaxation. Measurements of cell shortening and Ca transients in isolated mouse CMs. (f) Representative traces of sarcomere length (g) Fractional Shortening (h) Maximum rate of contraction (i) Maximum rate of relaxation (j) Representative traces of calcium transients (k) Calcium Amplitude (Fura-2 ratio-%baseline) (l) Maximum rate of Ca transient increase (m) Maximum rate of Ca transient decay. a-b: N = 6-8 mice per group. c-e: N = 6-7 mice per group. f-m: N = 6-7mice per group, n≥70 cells per group. All data are presented as mean ± SEM. P values were determined by unpaired, two-tailed, Student T test, *p ≤ 0.05, **p ≤ 0.01.

*In vivo* cardiac function was first assessed by echocardiography (Table S1). TgRRB mice had significantly greater LV fractional shortening and ejection fraction (Fig. 1b, Table S1). This occurred without difference in heart weight (Fig. S1), systolic ejection time (SET) and LV end-diastolic dimensions (Table S1) between TgRRB and WT hearts. The left-ventricular myocardial performance index (LV MPI), which reflects both systolic and diastolic function was significantly lower (indicating improved performance) in TgRRB mice as compared to WT. The gain in systolic function did not result in diastolic dysfunction as E/E’, a measure of LV filling pressure, and isovolumic relaxation time (IVRT), a measure of LV relaxation, were unchanged in the two groups (Table S1). Thus, the elevation of cytosolic [dATP] enhances systolic function without contributing to cardiac hypertrophy (Fig. S1) or impairing diastole (Fig. 1 and Table S1)[23].

To determine the effect of elevated dATP on cardiac function we used isolated Langendorff-perfused hearts to measure the *ex vivo* LV hemodynamics at baseline and with a high workload challenge. In paced hearts LV developed pressure (LVDP) product and both positive (+dP/dt) and negative (-dP/dt) rates of pressure development were greater for hearts from TgRRB vs WT mice (Fig. 1c, d, e). Interestingly, the rate-pressure product (RRP), a measure of cardiac output (Fig. S3b), was also greater for hearts from TgRRB mice. Myocardial responsiveness to high workload was elicited by elevating Ca^2+^. LVDP increased in both TgRRB and WT hearts, as did +dP/dt, in agreement with earlier findings that RNR overexpression does not compromise the heart’s ability to increase output[23,36] (Fig. S3d). Although mean LVDP and rate of pressure development were significantly higher, heart rate was slightly lower in unpaced TgRRB hearts (Fig.S3d, e). This resulted in similar rate pressure product between the two groups under higher workload conditions. Diastolic performance (-dP/dt) was comparable between the groups in response to the high-calcium challenge (Fig. S3h).

To determine the cellular basis of increased LV function, we measure single-cardiomyocyte (CM) Ca^2+^ transients and the resulting contractility. TgRRB-CMs exhibited significantly greater fractional shortening and maximum rates of contraction and relaxation as compared to WT cells (Fig. 1f). There was no significant difference in Ca^2+^ transient amplitude or time to peak amplitude in CMs from TgRRB versus WT mice and no change in baseline (resting) Ca^2+^ (Fig. 1j and Table S2). However, the rate of intracellular Ca^2+^ decay was greater in TgRRB-CMs. An increase in SERCA activity could explain the increased decay rate of the Ca^2+^ transient. However, there was no significant difference in sarcoplasmic reticulum Ca^2+^ store between the two groups. Thus, suggesting a different mechanism underlying faster Ca^2+^ decay observed in CMs with elevated dATP. In summary, all methods of increasing dATP enhanced contraction and relaxation in CMs.

Further, we examined the effect of chronic elevation of dATP on the sarcomeric protein function. We checked the phosphorylation state of sarcomeric protein - MyBP-C, Desmin, troponin T, troponin I, and MLC-2 in isolated myofibrils from hearts of both the groups (Fig. S4). There was no significant difference between the phosphorylation state of these sarcomeric protein which are possible modifiers of contractile properties in myocardium. Further, we studied Force pCa relationship and passive tension measurements in skinned cardiac muscle preparations (Fig. S6) from both the groups. Force generated at series of %stretch (sarcomere length) didn’t differ suggesting no difference in sarcomere elasticity within the groups (Fig. S6d)[14,37]. It is an indirect measurement of titin isoforms and its phosphorylation regulating the passive stiffness[38], suggesting no changes in the model of dATP chronic elevation. However, TgRRB trabeculae generated significant greater normalized force at pCa 4 than WT independent of dATP elevation (Fig. S6a, b). This suggest that even though we didn’t find difference in phosphorylation status of the sarcomeric protein, there is a possible myofilament adaption in myosin due chronic elevation of dATP, where more myosin heads interacting or interacting more strongly with actin even in absence of dATP elevation[20,22]. Nonetheless, TgRRB skinned trabeculae did respond to the dATP treatment like WT suggesting that the sensitivity to dATP is maintained in TgRRB trabeculae (Fig. S6d).

### Elevation of cardiomyocyte dATP leads to an increase in oxidative respiration capacity via increased myosin activation

In disease state of heart, there is a correlation between elevated crossbridge turnover rate and increased energetic cost of myocardial contraction[33]. Protein biochemistry measurements have shown this can also occur in resting cardiac muscle. The portion of myosin heads in sarcomeres in the disordered-relaxed (DRX) state (Fig. 3a) have a relatively high ATPase activity (~0.03 s^−1^) compared with the remainder of heads sequestered on the thick filament backbone in the super-relaxed (SRX) state with very low ATPase activity rates (~0.003 s^−1^)[39]. We recently reported that for myofibrils in solutions with 100% dATP, the SRX state is completely depopulated[40], resulting in significant increase in myofibrillar ATPase activity. However, it is unclear whether and how much ATPase in relaxed myofibrils is affected when the dATP content is ~1% of the total ATP pool, and how this may affect CM metabolism.

Thus, to determine how elevated dATP on the energetics of CMs, we measured oxygen consumption rate (OCR) of the TgRRB-CMs. These measurements were done on isolated cells cultured in assay medium with pyruvate and glucose following sequential addition of oligomycin (an inhibitor of ATP synthase), Carbonyl cyanide-p-trifluoromethoxy phenylhydrazone (FCCP, mitochondrial oxidative phosphorylation uncoupler), Antimycin A and Rotenone (mitochondrial electron transport chain inhibitor).

Basal respiration of TgRRB-CMs was approximately double that of cells from WT mice (Fig. 2c). TgRRB-CMs also had a 1.5-fold greater FCCP-stimulated maximal respiratory capacity (Fig. 2d). Subsequently, TgRRB-CMs show significantly elevated spare respiratory capacity (Fig. 2e). Spare respiratory capacity serves the increased energy demands when cells are subjected to stress, thereby maintaining cellular and organ function, cellular repair, or detoxification of reactive species[41]. Thus, TgRRB-CMs may perform better under high energy demands. Nevertheless, there was no significant difference between the groups in the extracellular acidification rate (ECAR), a surrogate for glycolytic activity (Fig. 2b). In CMs, factors such as cell size, sarcomere length, and beat rate can affect mechanical function as well as the oxygen flux and bioenergetics. The TgRRB-CMs did not exhibit differences in cell morphology or sarcomere length compared to WT-CMs (Fig. S1b).

**Fig. 2.**
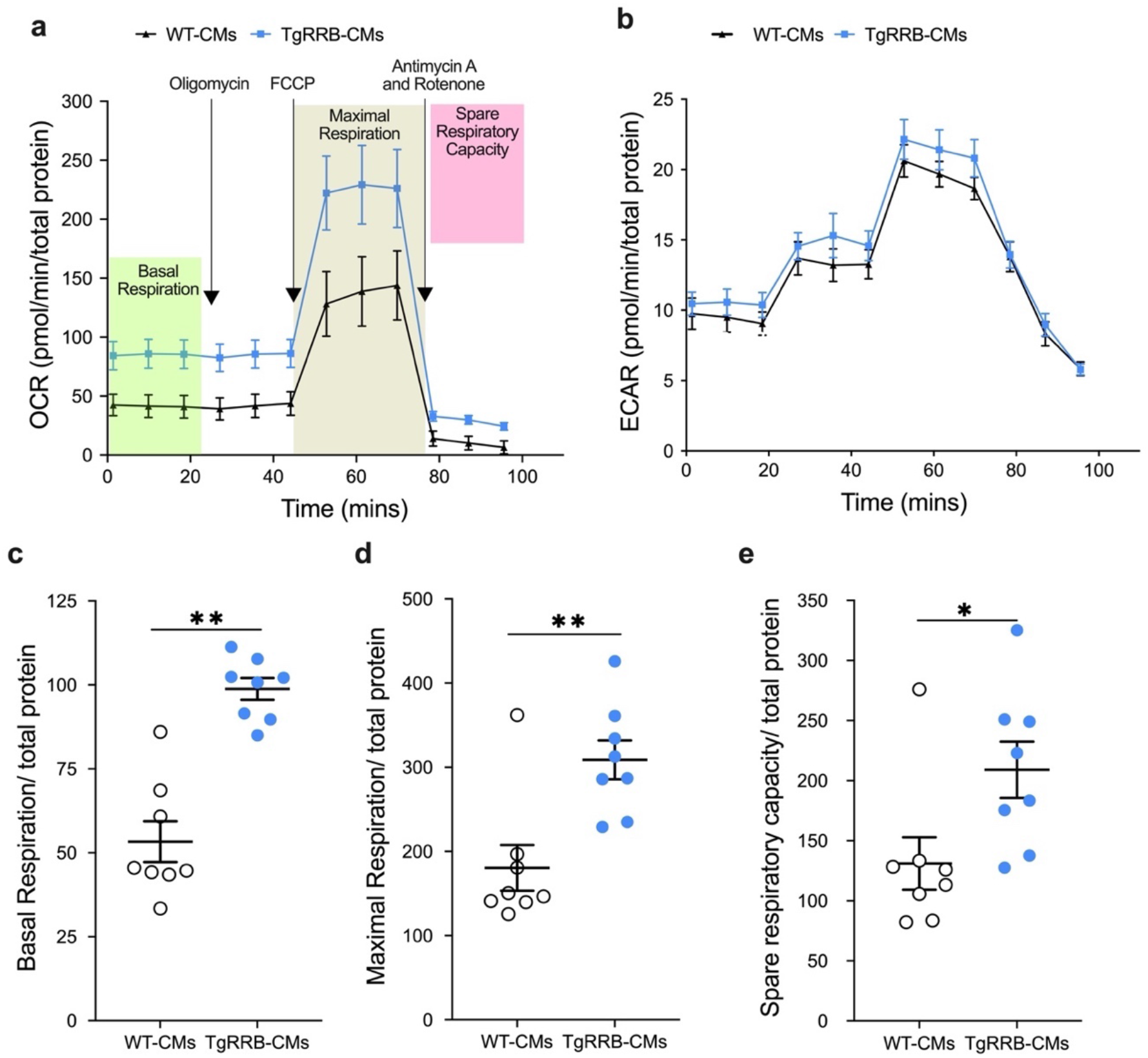
Assessment of cellular bioenergetics in TgRRB-CMs. Oxygen consumption rate (OCR) was measured on a standard Seahorse Mito stress test. (a) Representative OCR plot of CMs isolated from TgRRB vs WT mice (SD). (b) Representative ECAR plot of adult mice CMs isolated from TgRRB vs WT mice (SD). (c) Basal OCR (d) Maximal OCR and (e) Spare respiratory capacity normalized to the total protein. (c-d) N=8 mice per group, n≥3 technical replicates / animal. All data are presented as mean ± SEM. P values were determined by unpaired, two-tailed, Student T test, *P<0.05, **P<0.01.

Basal respiration is usually strongly controlled by ATP turnover via myosin ATPase, SERCA ATPase and other ion pumps and possibly be reflective of the higher cell shortening in TgRRB-CMs (Fig. 1f). Interestingly, our assessment of SERCA activity in mice cardiac tissue did not show a preference for ATP or dATP as substrate (Fig. S5) and we found no difference in sarcoplasmic reticulum Ca^2+^ load between TgRRB-CMs and WT-CMs (Table S2), suggesting that SERCA ATPase may not contribute to the basal respiration of TgRRB-CMs. Thus, these data suggest that the dATP elevation affects cellular bioenergetics, likely via modulation by myosin ATPase activity, as we have recently shown[21].

To address whether higher OCR was due to dATP mediated changes in the myosin SRX/DRX ratio[21], we treated TgRRB-CMs with Mavacamten (a selective myosin-motor inhibitor). Mavacamten is known to reduce myosin available for cross bridge cycling by recruiting myosin into the SRX state, leading to lower ATPase activity (Fig. 3)[42]. Acute treatment (1h) of Mavacamten reduced basal respiration of the TgRRB-CMs to WT-CMs levels (Fig. 3c) but did not affect the maximal respiratory capacity (Fig. 3d), thus supporting the idea that elevation in basal OCR is mainly due elevated myosin ATPase activity in TgRRB-CMs [40].

**Fig. 3.**
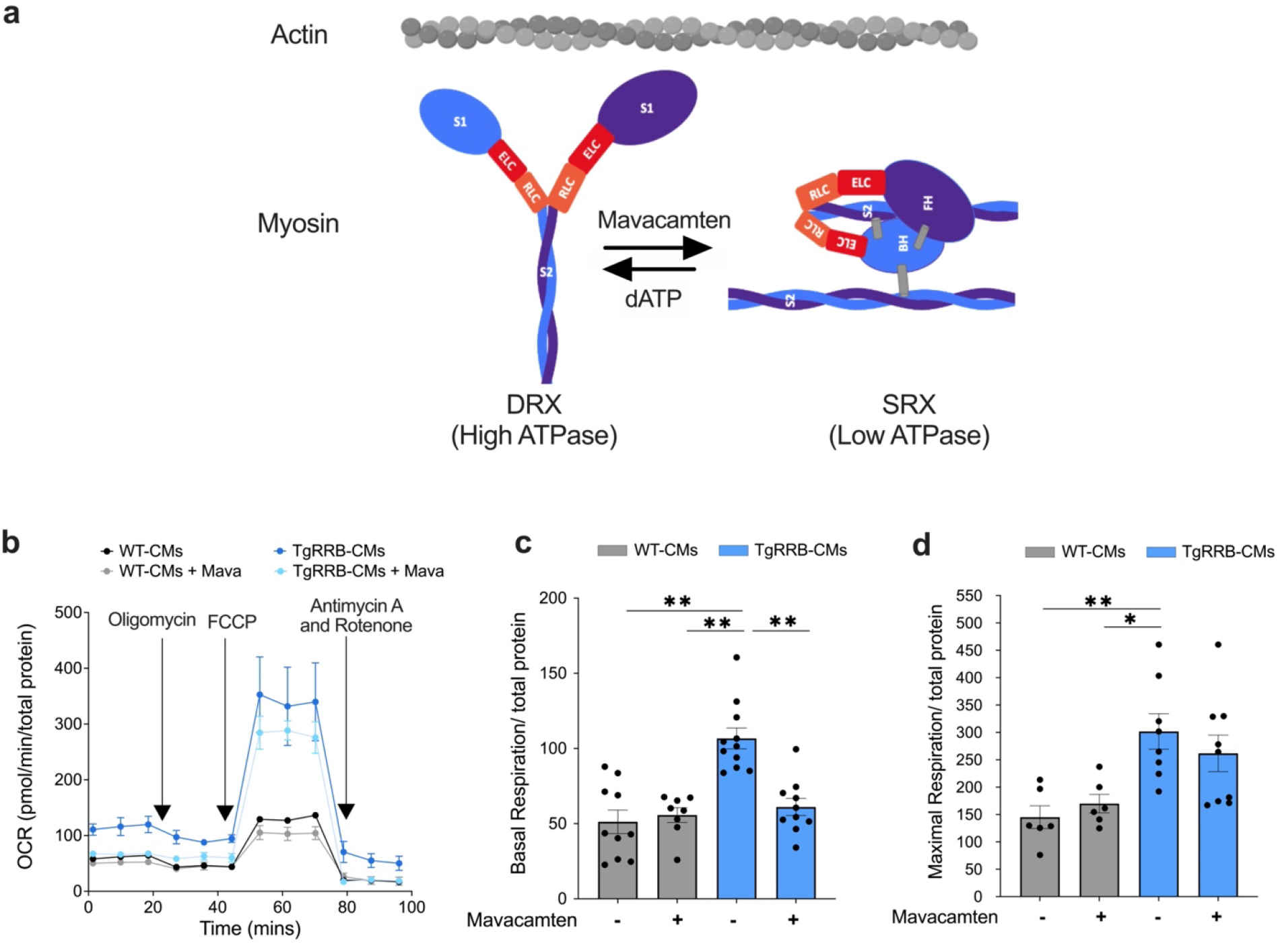
Constitutive dATP elevation leads to an increase in basal respiration via modulating myosin activation state. (a) The myosin ATPase states (DRX and SRX) can be modulated by Mavacamten and dATP[21]. ELC: essential light chain: RLC: regulatory light chain, BH: blocked (S2) head, FH: free (S2) head. Actin filament is shown in grey. Assessment of oxygen consumption rate (OCR) was performed with and without 1h treatment of Mavacamten (Mava, 3 μM) on adult mice CMs isolated from TGRRB vs WT mice (b-d). (b) Representative OCR plot, (c) Basal OCR (d) Maximal OCR normalized to the total protein. N=3 animals/group, n≥3 technical replicates / animal. All data are presented as mean ± SEM. P values were determined by ordinary one-way ANOVA with multiple comparison, *P<0.05, **P<0.01.

Importantly, in addition to basal respiration, mitochondria-dependent FCCP-induced -maximal oxygen flux was higher in TgRRB-CMs (Fig. 3d). The FCCP stimulated OCR suggests significant differences in the maximal mitochondrial capacity and electron transport chain (ETC) activity between TgRRB-CMs and WT-CMs, and these differences were not affected by Mavacamten. Thus, the mitochondria in TgRRB cells may have independent mechanisms of compensation (Fig. 2d and e) with elevation of dATP, improving spare respiratory capacity of these CMs.

### Metabolomics profiling reveals that TgRRB-CMs have elevated oxidative phosphorylation (OXPHOS) metabolite levels

We performed metabolomic analyses to understand the metabolic consequences of long-term elevation of dATP and the subsequent enhanced myosin activity. Targeted metabolomic analyses by LC-MS of isolated CMs from TgRRB mice and age-matched WT mice at 5-7 months of age measured 184 metabolites (Fig. S8). PCA of all 184 metabolites clearly separate the profiles of TgRRB-CMs (blue) from control WT-CMs (grey) (Fig. S8). The targeted analyses allowed assessment of altered pool sizes of intermediates relevant to glycolysis, the tricarboxylic acid cycle (TCA), fatty acid oxidation (FAO), and the pentose phosphate pathway (PPP). In the TgRRB-CMs there was a significant increase in pyruvate and all the TCA cycle intermediates, and a significant reduction in lactate/pyruvate ratio (Fig. 5a), suggesting a shift towards higher levels of pyruvate entering the TCA cycle. There were no differences in glycolytic intermediates (Fig. 4). Since there were lower levels of most PPP metabolites compared to WT-CMs, it suggests rerouting towards glucose oxidation in TgRRB-CMs. Amino acids such as phenylalanine, tryptophan, glutamate, serine, isoleucine, and glycine were more abundant in TgRRB-CMs compared to WT-CMs (Fig. 5d), and there was a trend toward the accumulation of citrulline, suggesting activation of the urea cycle and amino acid anaplerotic feeding of the TCA cycle. Thus, in TgRRB-CMs, there was an overall increase in OXPHOS compared to WT-CMs.

**Fig. 4.**
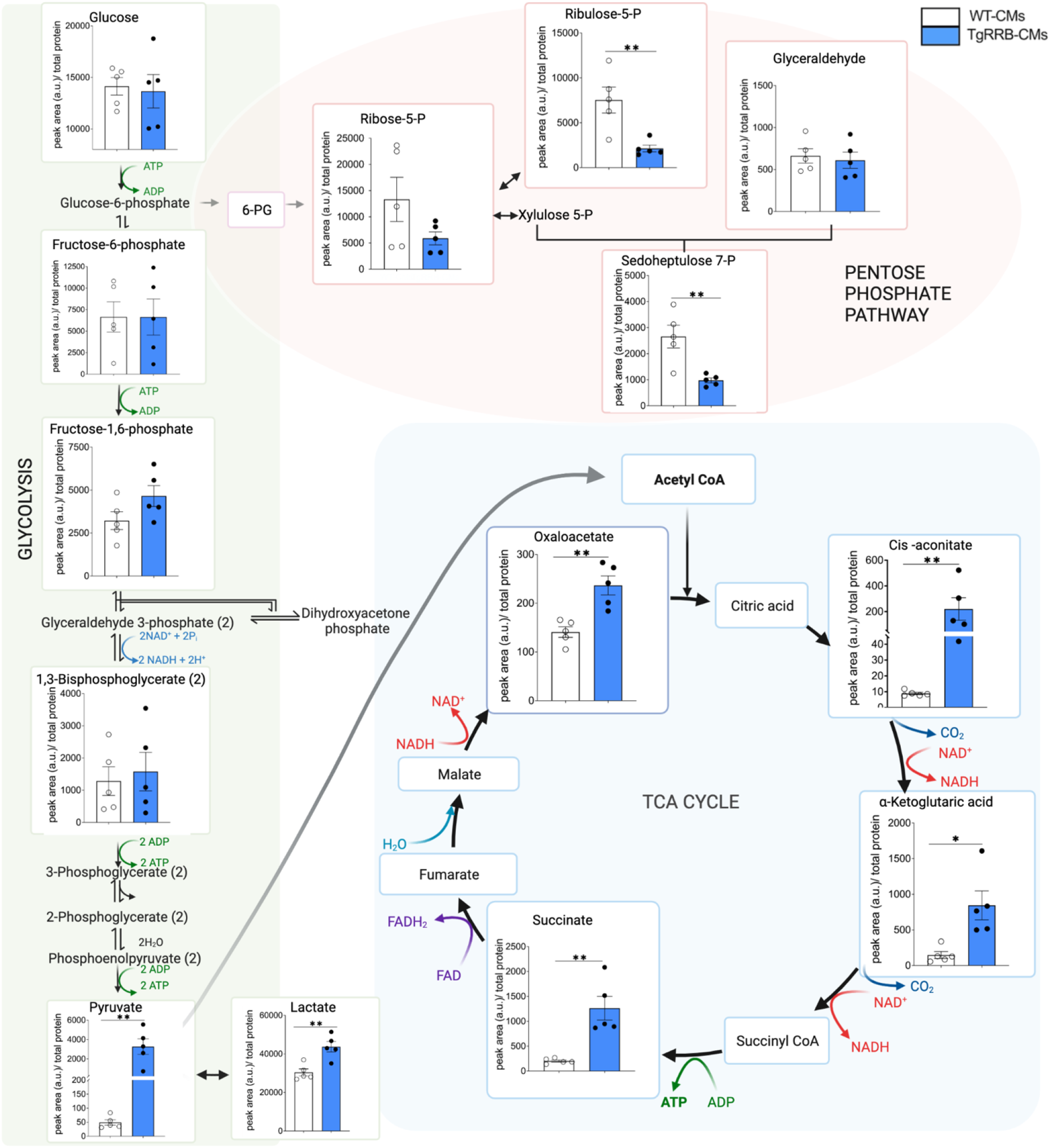
Pathway analysis of metabolites involved in Glycolysis, Pentose phosphate pathway, TCA cycle, in isolated CMs from TgRRB vs WT. Pathway analysis obtained from mass spectrometry-based metabolomics. N=5 animals/group. All data are presented as mean ± SEM. P values were determined by unpaired, two-tailed, Student T test, *P<0.05 **P<0.01. (Created with Biorender.com)

**Fig. 5.**
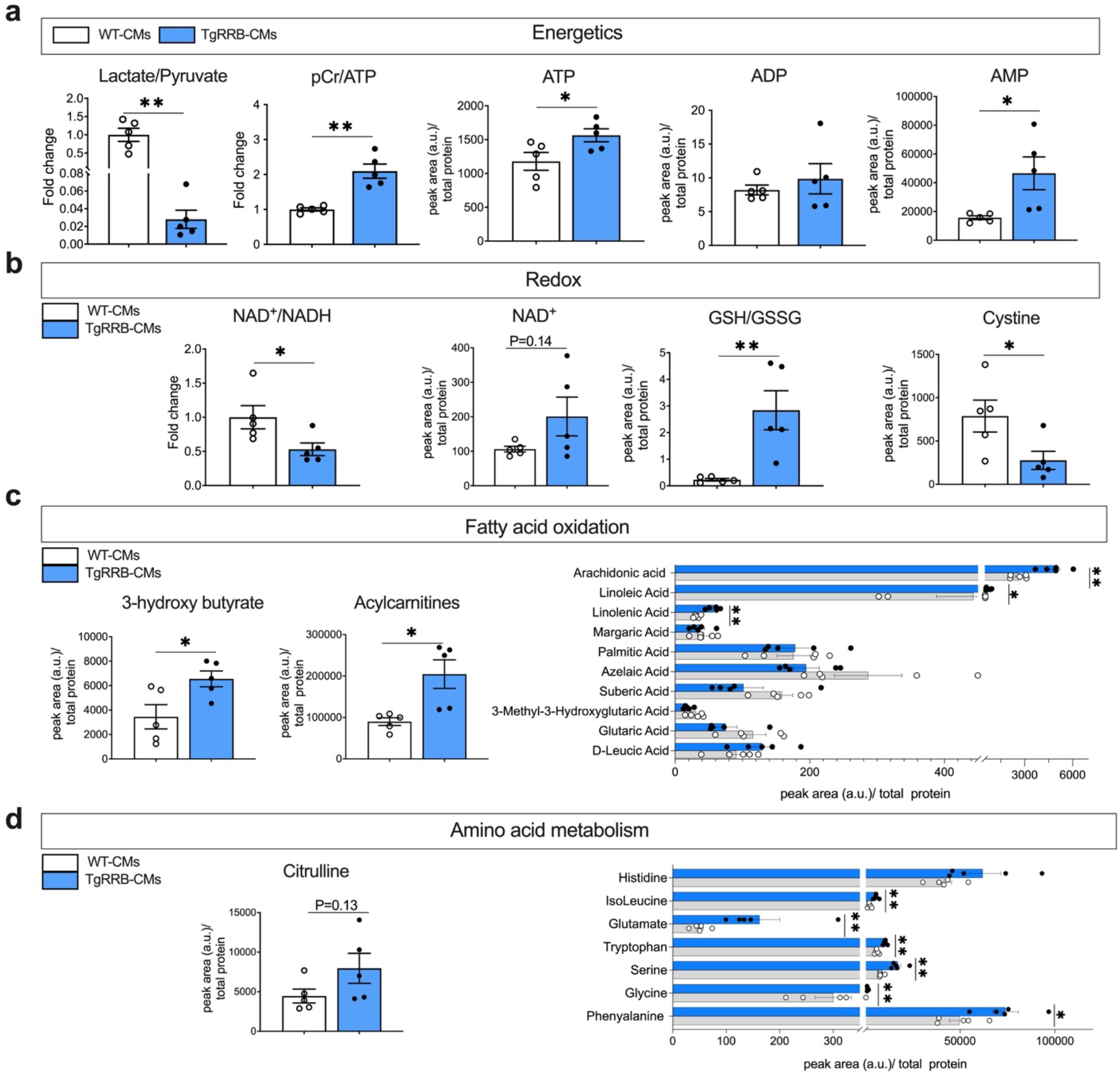
dATP elevation alters cellular metabolism and energetics. Mass spectrometry assessment of cellular metabolites involved in (a) Energetics: Lactate/Pyruvate fold change, phosphocreatine (pCr)/ATP fold change normalized to WT-CMs, ATP, ADP and AMP abundance in TgRRB -CMs vs WT-CMs (b) Redox: NAD^+^/NADH fold change, GSH/GSSG Fold change normalized to WT-CMs, NAD+ and Cystine abundance in TgRRB -CMs vs WT-CMs (c) Fatty acid oxidation: 3-hydroxy butyric acid, acylcarnitine and free fatty acids abundance in TgRRB -CMs vs WT-CMs (d) Amino acid metabolism: Citrulline and amino acids abundance in TgRRB -CMs vs WT-CMs. N= 5 animals per group. All data are presented as mean ± SEM. P values were determined by unpaired, two-tailed, Student T test, *P < 0.05 **P<0.01.

Stimulation of OXPHOS in the TgRRB-CMs is also evident from the NAD^+^/NADH and phosphocreatine (pCr) /ATP ratios as compared to WT-CMs (Fig. 5a, b). The latter was due the phosphocreatine levels being raised more than the modest increase in ATP content in transgenic CMs. Interestingly, there were also significantly greater AMP levels in TgRRB-CMs (Fig. 5a). Overall, therefore, TgRRB-CMs have an oxidative metabolic profile that supports the observed higher respiration rate.

In healthy myocardium, oxidative stress via OXPHOS is tightly counterbalanced by reducing agents. In TgRRB-CMs, we found that the metabolites involved in the endogenous antioxidant defense mechanism, such as lactate and glutathione (GSH), were significantly higher (Fig. 5b). Oxidized cysteine (cystine), a marker for oxidative stress, was reduced. Thus, greater levels of protective antioxidant metabolites were found in TgRRB-CMs.

TgRRB-CMs also appear to have increased fatty acid metabolism, with greater levels of 3-hydroxybutyric acid (more than 1.5-fold) and intermediate products of oxidative decomposition of fatty acids (Fig. 5c). These findings are suggestive of increased ketone body metabolism and FAO in the TgRRB versus WT-CMs. Compared to WT-CMs, TgRRB-CMs were also enriched with acylcarnitine and free fatty acids (especially polyunsaturated fatty acids like linolenic and arachidonic acids), consistent with an elevated beta-oxidative capacity of TgRRB-CMs to utilize fatty acids as an energy source (Fig. 5c).

Overall, the metabolomic analysis suggests that TgRRB-CMs exhibit a significant metabolic shift towards increased oxidative metabolism, combined with greater phosphocreatine kinase and high energy phosphates, ultimately improving the energetics of TgRRB-CMs (Fig. 4, 5).

### TgRRB-CMs have enhanced mitochondrial function compared to WT-CMs

We next sought to characterize whether the metabolic and energetic differences in TgRRB-CMs versus WT-CMs were associated with changes in cardiac mitochondria. Increased maximal respiration of TgRRB-CMs could occur via increased mitochondria biogenesis or increased activity of existing respiratory chain proteins. To explore this, we studied regulators of cell energetics, mitochondrial content, morphology, and function (Fig. 6).

**Fig. 6.**
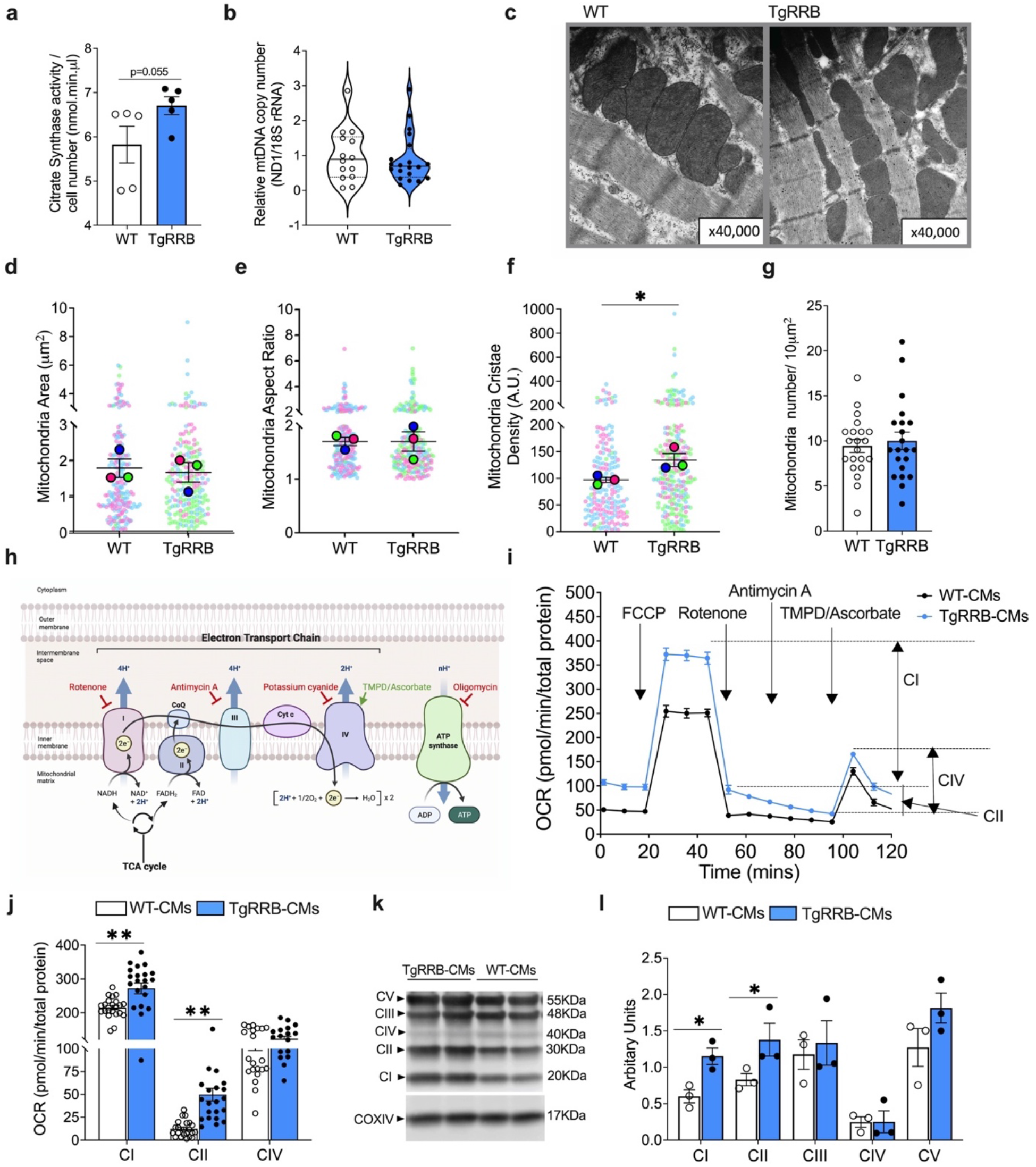
The mitochondrial structure and function in TgRRB-CMs and WT-CMs. (a) Citrate synthase (CS) activity (N=5 animals/ group) (b) Relative mitochondrial DNA copy number. Mitochondrial DNA (ND1; mitochondria-encoded NADH dehydrogenase 1 gene; mtDNA) was measured relative to 18S rRNA (nucleus-encoded gene; nDNA) as the control (c) Representative electron photomicrographs of mitochondria at a magnification of x40,000.

The expression analysis of genes involved in the regulation of mitochondria biogenesis and cell energetics, such as PGC1β (peroxisome-proliferator–activated receptor-γ-coactivator 1 β), TFAM (transcription factor A were greater in TgRRB hearts suggesting that there is greater OXPHOS gene expression[43,44] (Fig. S10). Citrate synthase activity analysis showed a trend for higher levels in TgRRB-CMs compared to WT-CMs (Fig. 6a). Nonetheless, mitochondrial content, determined via the mtDNA to nuclear DNA ratio, was not significantly different (Fig. 6b). We next examined mitochondrial ultrastructure using transmission electron microscopy. Mitochondrial number, size, aspect ratio were not different from WT group (Fig. 6d, e, g). However, mitochondrial cristae density was significantly higher (38.4 % ± 0.14) in TgRRB heart tissue as compared to WT, suggestive of an enhancement in the ETC complex situated on cristae (Fig. 6f). Overall, these measurement suggest a potential increase of mitochondrial turnover activity and quality control in TgRRB hearts[43].

A relatively higher ETC content may be elevating the oxidative capacity of the mitochondria in TgRRB-CMs. We examined the overall mitochondrial activity and function of the electron transport chain (ETC) proteins in TgRRB-CMs. A standard Seahorse Mito stress test was used to measure oxygen flux in response to ETC complex inhibitors in sequence, to parse out complex-(es) whose activity was affected (Fig. 6h, i)[45]. TgRRB-CMs had significantly greater CI and CII activities compared to WT-CMs (Fig. 6j). We further confirmed higher levels of complex I (CI) and II (CII) proteins in TgRRB-CMs compared to WT-CMs (Fig. 6k, l). Together, these results suggest that chronic elevation of dATP improves mitochondrial respiration via elevated expression and activity of CI and CII activity, fueled by elevated OXPHOS (NADH and FADH2) in the TgRRB versus WT-CMs.

Scatter plot showing morphometric analysis (X40,000) of sarcomeric (d) mitochondria area, (e) aspect ratio, (f) cristae density with solid symbols as average per animal. X8000 images were analyzed for (g) average mitochondria number per 10μm^2^ per animal. (h) Schematic of different compounds targeting Complex I (CI), Complex III (CIII), Complex IV (CIV) and ATP synthase. (i) Representative OCR plot with the application of compounds to assess ETC complex activities of mitochondria. (j) Bar graph depicting C I, Complex II (CII), and CIV activities (k) Western blot of mitochondrial ETC complex proteins and (l) densitometry analysis normalized to COXIV levels. N=3 animals/ group. All data shown in mean ± SEM. P values were determined by unpaired, two-tailed, Student T test, *P<0.05, **P<0.01.

## Discussion

We have previously demonstrated that dATP is a candidate myosin activator for heart failure therapy and established that it can improve contraction and relaxation in a variety of heart failure models[46,47]. However, an important question that needed to be addressed is whether or how elevation of dATP affects cardiomyocyte (CM) metabolism. The present study demonstrates that a modest elevation of dATP levels in CMs that is sufficient to improve LV and CM function, also improves metabolism and cellular bioenergetics, providing crucial insights into the relationship between energy supply-demand and crossbridge activity in CMs.

We employed novel strategies to stably overexpress RNR, utilizing ubiquitylation-resistant variant RRM2B in a transgenic mice model (TgRRB) and found that dATP level was significantly elevated from <0.1% to ~ 0.5-1% (of the total ATP pool) in TgRRB mouse heart (Fig. 1a, S2). This novel strategy yielded important insights. First, it appears that this approach, to elevate dATP with a proteolytically resistant variant RNR is limited to dATP content of ~1% of the total ATP pool[48]. This relatively low-level elevation of dATP is sufficient to improve systolic as well as diastolic function in the heart *in vivo* and in isolated CMs *in vitro*. Second, we demonstrated that cells with elevated dATP have elevated basal oxygen consumption due to changes in myosin state, independent of active crossbridge cycling. Third, these cells respond with a metabolic shift, such that there are higher pools of oxidative metabolites, with elevated OXPHOS, FAO, and energy generation in TgRRB-CMs. Finally, we provide evidence that the long-term mitochondrial remodeling in CMs may occur to accommodate for the higher energy demands of the high functioning dATP CMs.

The current study demonstrates that long-term elevation of cardiac [dATP] results in sustained elevation of basal left ventricular performance while maintaining high workload and energetic reserve, with no signs of hypertrophy or altered diastolic dimensions[23]. This is important because in some cardiomyopathies with elevated workload, there is cellular hypertrophy along with other pathological remodeling. The improved twitch function with no significant changes in Ca^2+^ transient amplitude (Fig.1j and Table S2, S3), suggest differences in the cellular contractility are due to dATP-mediated activation of myosin heads[20]. Interestingly, the skinned myocardial force measurements revealed that there might be adaptation in the myosin structure and function with chronic exposure to dATP elevation in TgRRB (Fig. S6). Nonetheless, TgRRB samples did remain sensitive to dATP elevation leading to increased force generation. Cellular relaxation kinetics are also improved, perhaps due to improved calcium decay kinetics (Fig. 1m) and the maintenance of the myosin in its activated state during relaxation, which may further facilitate subsequent contractions[22,49]. <u>Thus, dATP increases myosin cycling and sarcomere function without prolonging the systole (Fig.1, Table S2)</u>[47,50] <u>or shortening diastole, which possibly makes it an energy-efficient myotrope.</u>

In CMs, myosin is the primary mechanoenzyme coupling the energy released from ATP hydrolysis to force generation. Biochemical studies have shown that myosin heads (S1, Fig. 3a) are present in multiple physiological/conformational states in resting muscle, and these have different rates of ATP consumption which, in turn, when modulated (pathological mutation) can affect the overall CM metabolism[32,51]. Our recent study demonstrated that biochemically dATP pushes myosin heads from a low ATP consumption (SRX) to higher ATP consumption (DRX) state (Fig. 3a)[21]. Here we demonstrated that the dATP induced change in SRX/DRX underlies the greater basal respiration of TgRRB-CMs (Fig. 3c). Thus, demonstrating that the effect of dATP on basal energetics is by myosin state modulation.

In the present study, we also found that dATP elevation rewires cellular metabolism with to avoid energy deficit and downstream pathology. Metabolomic analysis showed that dATP elevation in TgRRB-CMs results higher OXPHOS with greater levels of pyruvate, FAO metabolites (Fig 5c), TCA intermediates (Fig. 4), and anaplerotic metabolites (Fig. 5d), but not in the levels of PPP metabolites (Fig. 4). The shift away from PPP to glycolysis might add to the pyruvate accumulation. Alternatively, FAO may be responsible for filling up the TCA cycle metabolites such that glycolysis needed is less and can cause pyruvate to build up. Overall, TgRRb-CMs had a modest but significant increase in ATP levels (Fig. 5a). Interestingly, we also found elevated levels of reducing agents (e.g., GSH/GSSG, NADH) indicating low oxidative stress in high energy TgRRB-CMs (Fig. 5b).

The oxidative metabolic profile of the TgRRB-CMs is supported by high functioning mitochondria displaying higher cristae density and number of ETC complexes and their activity in TgRRB (Fig. 6f, j, l), indicating an improvement in mitochondrial intrinsic ATP synthesis capacity in TgRRB-CMs[52]. This adaptation may be the consequence of a trade-off between the elevated mechanical work (contraction), sarcomere architecture, and oxidative metabolic power (OCR) as produced by mitochondria and serves as a mechanism for increasing metabolic power without constraining contractile function. Thus, TgRRB-CMs show shift in metabolism towards OXPHOS and high mitochondrial oxidative capacity to generate ATP, thereby avoiding activation of anaerobic respiration observed in pathological condition[31,32].

In this manner, dATP mediated steady elevated cellular respiration does not compensate LV contractile function– e.g., normalize pressure to WT levels – but remodels cellular metabolism in the long run via more oxidative metabolism and increased mitochondrial turnover and ETC activity. We showed that dATP-induced myosin activation elevates resting metabolism and contractility in CMs; however, the mechanism behind how this elevated flux induces enhanced OXPHOS and mitochondrial remodeling is unknown. TgRRb mice were lower in body weight (Fig. S1) than WT littermates suggesting that the whole-body metabolic rate might be affected. Intriguingly, dATP can be used as a tool to understand how cellular as well as body metabolism responds to increased myosin cross bridge activity. Overall, TgRRB mice demonstrate beneficial compensatory changes in CM metabolism to accommodate for the gain in ventricular function.

In summary, we provide the first evidence that a small increase (~1%) dATP via persistent overexpression of RNR variant promotes cardiac contractile function putatively by elevating basal ATPase activity of myosin and adapts cellular metabolism towards higher oxidative capacity. Elevated cellular and mitochondrial dATP (Fig. S2) has a panoply of beneficial effects on cardiac metabolism, from remodeling of cellular metabolic pathways and respiration to mitochondrial cristae alterations and function representing dATP as a potential therapeutic option for targeting the contractile and metabolic deficit observed in heart diseases.

### Study limitations

A limitation of the study is its dependence on isolated Langendorff-perfused hearts for assessing cardiac function. Although isolated Langendorff-perfused heart preparation provides a way to investigate heart function independent of the neurohumoral and physiological adaptations, other physiological measurements can give more insights into the effect of elevated dATP on other factors regulating cardiac function, like afterload. With long-term exposure to dATP in this model, we did observe dATP-independent differences in force generation, possibly indicating secondary effects of chronic elevation of contractility. Seahorse assay and metabolomics in this study were performed on isolated CMs from TgRRB mice and age-matched WT mice. This approach has the clear advantage of exclusively looking at the effect of myotrope dATP on CM metabolites and energetics. However, it needs to be specified that metabolism could be altered during CM isolation and time in culture during the assay. <u>Also, seahorse measurements are carried out on unloaded cells, which could affect the interpretation of the energy consumption. Despite this limitation, we observed significant differences in the metabolism and energetics of TgRRB-CMs and WT-CMs, emphasizing the independent metabolic effects of dATP. Since all the measurements carried out in this study are in a healthy heart, a critical next step is to study the metabolic consequence of dATP elevation in a stressed heart. This enhanced understanding of the role of metabolic regulation in the heart via dATP should open possibilities for harnessing these processes for therapeutic potential.</u>

## Materials and Methods

All animal experiments were approved by the University of Washington (UW) Animal Care Committee. Animals were cared for in accordance with the US National Institutes of Health Policy on Humane Care and Use of Laboratory Animals in the Department of Comparative Medicine at UW.

### Echocardiography

Echocardiography was performed on mice at four months of age (Vevo 3100, Fujifilm Visual Sonics, Toronto, Canada) as previously described [23]. The parasternal short axis view at the mid-papillary level was used to obtain M-mode images for analysis of LV end-diastolic (LVEDD), end-systolic (LVESD) dimensions, fraction shortening (SF) and ejection fraction (EF). Diastolic function was assessed by measurement of trans mitral flow parameters, including systolic ejection time (SET) and isovolumic relaxation time (IVRT) from apical 4-chamber view with pulsed wave Doppler. Tissue Doppler imaging was performed by placing sample volume at the septal corner of the mitral annulus to measure early diastolic myocardial relaxation velocity (E’). All analysis was done by a blinded single reader.

### Isolated perfused mouse heart preparation

Left ventricular contractile kinetics were measured in Langendorff isolated heart preparations as described previously [36]. Excised mouse hearts were perfused at a constant pressure of 80 mmHg in a modified Krebs-Henseleit buffer (In mM: 118 NaCl, 25 NaHCO_3_, 5.3 KCl, 2.0 CaCl_2_, 1.2 MgSO_4_, 0.5 EDTA, 5.5 glucose, 0.5 sodium pyruvate, equilibrated with 95% O_2_ and 5% CO_2_, pH 7.4) at 37°C. After equilibration, baseline function was monitored at a fixed end-diastolic pressure of 8-10 mmHg for 10-20 minutes by way of a water-filled balloon inserted into the LV. Balloon volume was increased in 5 μL increments to determine the pressure-volume relationship in the myocardium. Finally, buffer [Ca^2+^] was increased from 2mM to 4mM for 20 minutes to stimulate an increase in cardiac work.

### Adult mouse cardiomyocyte isolation and contractile assessment

CMs were isolated from mice via enzymatic digestion through retrograde aortic perfusion as described previously [23]. Briefly, hearts were rapidly excised, cannulated, and perfused with a collagenase/protease solution, following which the ventricles were removed, minced, and triturated. Contractile assessments of cells were made immediately after stepwise reintroduction of calcium and incubation in Fura-2-AM (Invitrogen) to measure calcium transients. Cell shortening and re-lengthening and calcium transients were measured at 1 Hz and 37°C using an IonOptix μStep video recording system (IonOptix, Westwood, MA, USA). Measurements were performed by four independent experimentalists and analyzed by the same reader. In addition, we also measured Sarcoplasmic Reticulum (SR) Ca^2+^ load using the caffeine technique. Briefly, isolated CMs will be loaded with Fluo-4 Ca^2+^ indicator dye, then paced (1 Hz for 8-10s) to stabilize Ca^2+^ transient amplitudes before pico-spritzing a 20 mM caffeine solution.

### dATP quantification in cardiac tissue

dATP content in CMs was measured by high-performance liquid chromatography-mass spectrometry (HPLC-MS/MS) as described previously [53]. Freshly isolated whole mouse ventricle or CMs were flash-frozen, stored at −80°C, and nucleotides were extracted 1-3 days before measurement using a 50% methanol solution. The supernatant was stored at −20°C until ready for injection into the HPLC-MS/MS system. ATP and dNTP standard curves were used to calculate the concentration of each nucleotide. dATP content was then normalized to ATP in the same samples.

### Western blot

Protein samples were solubilized in 1% NP-40 lysis buffer with 4% protease inhibitors (Sigma) and centrifuged for 14,000 g for 5 minutes. The supernatant was mixed with SDS protein sample buffer containing β-mercaptoethanol (Bio-Rad) and resolved by SDS-PAGE on 4-20% Mini-Protean GTX Stain-Free gels (Bio-Rad). Protein transfer to 0.45 mm polyvinylidene difluoride membranes was performed at constant 100 volts at 4°C in Towbin’s buffer containing 20% methanol for 1 hour. Blots were blocked for 1 hour at room temperature in 1x BlockerFL (Thermo Scientific) before overnight incubation with primary antibodies (Total OXPHOS Rodent WB Antibody Cocktail: Abcam ab110413, 1:500; Anti-COX IV antibody - Mitochondrial Loading Control: Abcam ab16056, 1:500; Human RRM1: Abcam ab137114, 1:500; Human RRM2/RRM2B: ThermoFisher PA579937, 1:1000 Human Rrm2B: ThermoFisher MA5-29529 1:1000, GAPDH: Fitzgerald 10R-2932 1:1000). One-hour incubation with StarBright fluorescent secondary antibody (#12005866, #12004162 BIO-RAD) diluted 1:2500 in PBS-Tween 5% milk was used to detect primary antibodies bound to the blot by BIO-RAD ChemiDoc imaging system. The densitometry of western blots was performed using I Image Lab (BIO-RAD USA).

### Enzyme Assays

Citrate synthase activity, a key enzyme in the Kreb’s cycle that is commonly used marker for mitochondrial mass, was measured on tissue homogenates and isolated CMs with a kit purchased from Sigma-Aldrich (MAK193; St. Louis, MO, USA). In brief, CS activity is determined using a coupled enzyme reaction, which results in a colorimetric (412 nm) product proportional to the enzymatic activity present. We normalized the CS activity to the number of cells used for the assay.

### Metabolomic analysis

CMs samples were isolated from hearts of age matched (5-7 months) TgRRB and WT mice. The CMs were either plated for XFe24 bioenergetic profiling or flash frozen for Metabolomic analysis. Aqueous metabolites for targeted LC-MS analysis were extracted using a protein precipitation method. Samples were first homogenized in 200 μL purified deionized water at 4 °C, and then 800 μL of methanol containing 6C13-glucose and 2C13-glutamate (reference internal standards) was added. Afterwards, samples were vortexed, stored for 30 minutes at −20 °C, sonicated in an ice bath for 10 minutes, centrifuged for 15 min at 14,000 rpm and 4 °C, and then 600 μL of supernatant was collected from each sample. Lastly, recovered supernatants were dried on a SpeedVac and reconstituted in 1 mL of LC-matching solvent containing 2C13-tyrosine and 3C13-lactate (reference internal standards). Protein pallets that were left over from the sample prep were saved for the BCA protein assay.

Targeted LC-MS metabolite analysis was performed on a duplex-LC-MS system composed of two Shimadzu UPLC pumps, CTC Analytics PAL HTC-xt temperature-controlled auto-sampler and AB Sciex 6500+ Triple Quadrupole MS equipped with ESI ionization source. UPLC pumps were connected to the auto-sampler in parallel and were able to perform two chromatography separations independently from each other. Each sample was injected twice on two identical analytical columns (Waters XBridge BEH Amide XP) performing separations in hydrophilic interaction liquid chromatography (HILIC) mode. While one column was performing separation and MS data acquisition in ESI+ ionization mode, the other column was getting equilibrated for sample injection, chromatography separation and MS data acquisition in ESI-mode. Each chromatography separation was 18 minutes (total analysis time per sample was 36 minutes). MS data acquisition was performed in multiple-reaction-monitoring (MRM) mode. LC-MS system was controlled using AB Sciex Analyst 1.6.3 software. Measured MS peaks were integrated using AB Sciex MultiQuant 3.0.3 software. 205 metabolites and 4 reference standards were measured in the study samples. In addition to the study samples, two sets of quality control (QC) samples were used to monitor the assay performance as well as data reproducibility. One QC [QC(I)] was a pooled human serum sample used to monitor system performance, and the other QC [QC(S)] was pooled study samples, and this QC was used to monitor data reproducibility. The data were well reproducible, with average and median CVs of 5.2% and 4.3%, respectively. Generated MS data were normalized vs BCA total protein count. The PCA and pathway impact analysis of the metabolomic data were carried out using MetaboAnalyst 5.0 software. The fold change calculated for the metabolite ratios (Lactate/Pyruvate, phosphocreatine (pCr)/ATP, NAD^+^/NADH, and GSH/GSSG) are normalized to the WT group. Differences in the abundance of individual metabolites were determined using a student’s t test (two-tailed, unpaired comparison). A value of P < 0.05 was considered statistically significant. Data are given as mean ± SEM.

### XFe24 bioenergetic profiling

Isolated CMs were seeded on matrix gel–coated cell culture microplates (Agilent Seahorse XF24) at the cell intensity of 20000 cells/well. Generally, before performing the experiment, the culture medium was changed into 625uL assay medium (Agilent Seahorse XF Base Medium), and then, cells were incubated in 37°C non-CO2 incubator for 1 hour.

The bioenergetic response of cardiomyocytes (CMs) was measured with the Seahorse Bioscience XFe24 Flux Analyzer. For oxygen consumption rate (OCR) measurements, the plating media were changed to 675 μl DMEM-based Seahorse (Agilent, 102353-100) supplemented with 4 mmol/l glutamine, 1 mmol/l pyruvate and 5 mmol/l glucose 1 h before assay. The XFe24 automated protocol consisted of 10 min delay following microplate insertion, baseline OCR measurements [3 times (3 min mix, 2 min wait, 3 min measure)], followed by injection of port A (75 μl). The same protocol is followed for the port B (75 μl) and port C (75 μl). Optimal concentrations of oligomycin (Sigma, 75351), carbonyl cyanide-p-trifluoromethoxyphenylhydrazone (FCCP; Sigma, C2920), Rotenone (Sigma, R8875) and antimycin A (Sigma, A6874) were diluted in DMSO (Sigma, 154938). XF measures the oxygen consumption rate (OCR), represents mitochondrial respiration. The OCR measurements were normalized to the protein concentration. Experimental replicates were normalized to protein concentration. Optimization of seeding density was conducted in compliance with the Seahorse XF24 manufacturer recommendations through multiple experiments to identify a cell count that provided basal OCR readings sufficiently above background, within tested linear response ranges and with full re-oxygenation of the media attained between measurements.

### Transmission Electron Microscopy

Tissue samples were fixed overnight in ½ strength Karnovsky’s (2%paraformaldehyde/2.5% glutaraldehyde buffered with 0.2M cacodylate) and post fixed in 2% OsO4 buffered in 0.2M cacodylate buffer. After dehydration, tissues were embedded in Epon 812 (Electron Microscopy Sciences), thin sectioned (70 nm), and stained with uranyl acetate for 2 h and lead citrate for 5 min. The samples were imaged using a JEOL 1230 transmission electron microscope set to 80 kV, and the images were captured using a Gatan digital imaging system. n≥4 scans were taken per animals per magnification. Mitochondrial objects in the images were pre-segmented in an automated manner using ImageJ software: after noise reduction by repetitively convolving each image with a Gaussian function, images were thresholded by the ‘Percentile’ auto thresholding method. Boundaries between background and foreground were smoothened by repeated erosion and dilation. Lastly, particles were identified based on set cut offs for size and circularity. Results from this initial machine-based analysis were subsequently manually verified as mitochondrial particles in a blinded fashion. Image coordinates of each identified particle were used for the subsequent calculation of mitochondrial number, size, aspect ratio, while brightness provided an estimate for cristae density as described in [54].

### Statistics

The statistical analyses were performed using GraphPad Prism 8 (Graphpad Software). Results were considered statistically significant at p ≤ 0.05. The results shown here are mean ± SEM unless otherwise stated.

## Acknowledgements

We acknowledge Dr. Kristina B. Kooiker for her guidance and assistance with trabeculae force measurement and sarcomeric protein phosphorylation protocol. KM was funded by the Institute for Stem Cells and Regenerative Medicine fellows’ program. MR acknowledges NIH R01 HL128368, P30 AR074990 and, DOD MD170031. FMH acknowledges NIH Grant K08 HL128826 and the Locke Charitable Trust.

