## Supplementary Appendix for "dATP Elevation Induces Myocardial Metabolic Remodeling to Support Improved Cardiac Function"

#### Figures

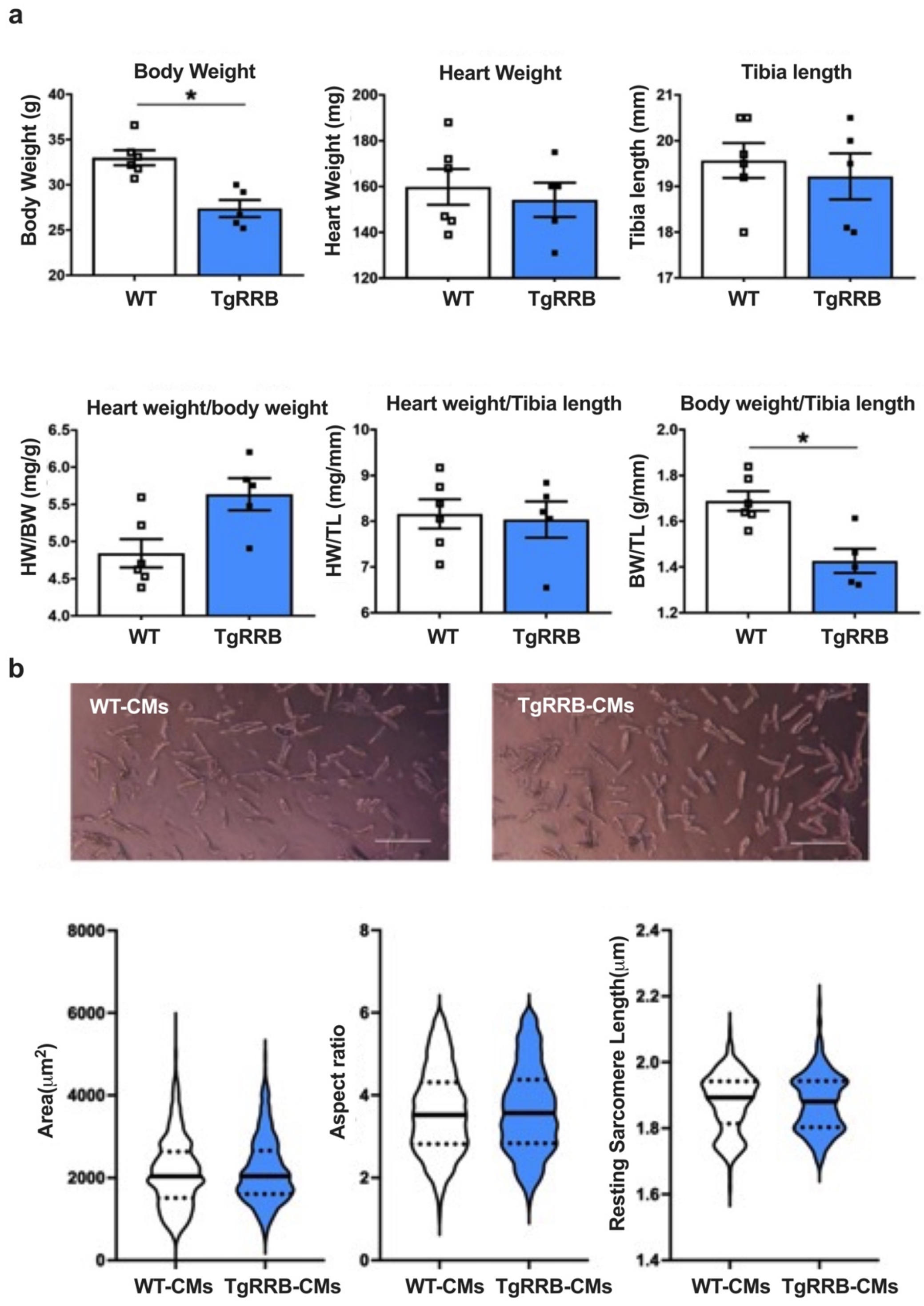

**Fig. S1. TgRRB mice were smaller in body weight with no cellular and gross cardiac hypertrophy.** (a) Body Weight, Heart weight and tibia weight (b) Representative bright field image of TgRRB-CMs than WT-CMs (scale bar: 150μm). Cellular area, aspect ratio and resting sarcomere length measurements. N = 5-6 mice per group. All data are presented as mean ± S.E.M. P values were determined using unpaired, two-tailed, Student's t-tests. *p ≤ 0.05.

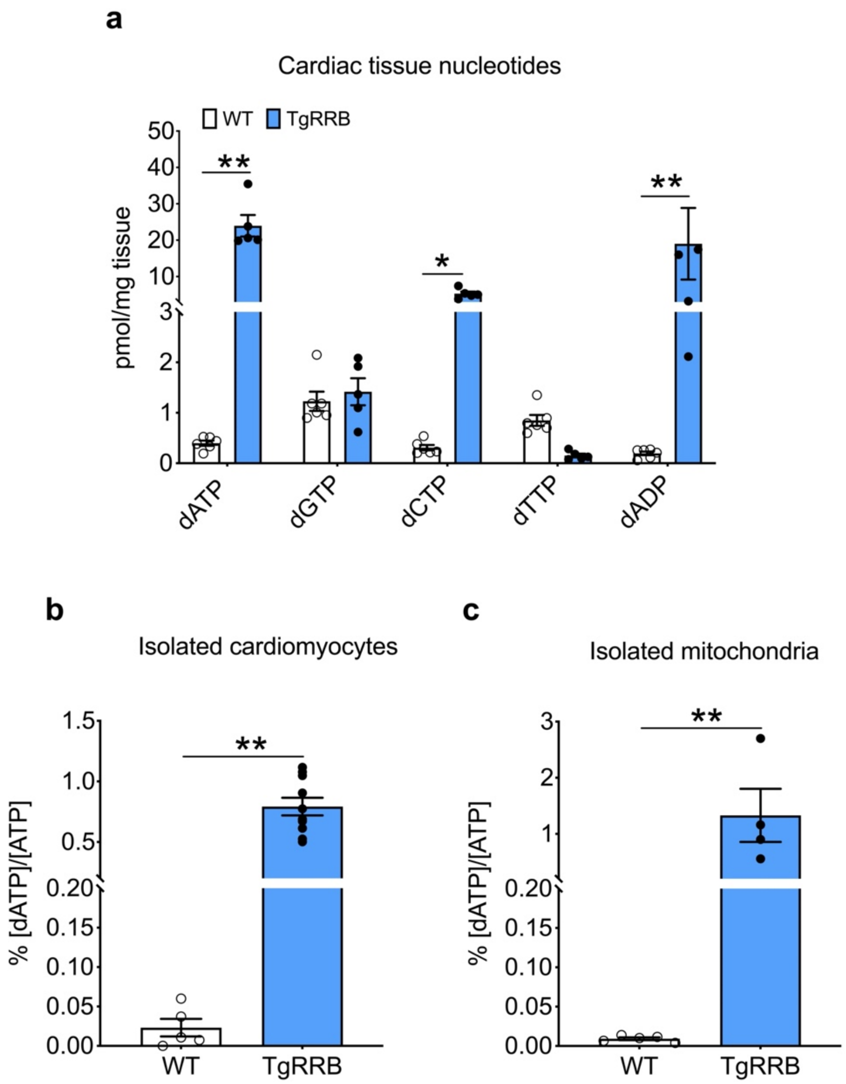


**Fig. S2. Nucleotide levels in TgRRB vs WT.** Mass spectrometry measurements of deoxynucleotides in (a) heart tissues. dATP/ATP% in (b) isolated CMs and (c) isolated mitochondria. N=4-6 mice per group. All data are presented as mean ± S.E.M. P values were determined using unpaired, two-tailed, Student's t-tests. **p ≤ 0.01.

| Systolic measurements | WT | TgRRB |
|---|---|---|
| HR, bpm | 514.81±42 | 531.35±38 |
| LVID; s, mm | 2.38±0.17 | 2.21±0.14** |
| LVID; d, mm | 3.86±0.18 | 3.85±0.17 |
| LVAW; s mm | 1.57±0.23 | 1.69±0.16 |
| LVPW; s mm | 1.30±0.11 | 1.32±0.12 |
| LVAW; d mm | 1.05±0.217 | 1.08±0.1 |
| LVPW; d mm | 0.94±0.08 | 0.89±0.07 |
| EF, % | 69.01±2.7 | 74.25±2.43** |
| FS, % | 38.3±2.03 | 42.60±2.15** |

| Diastolic measurements | WT | TgRRB |
|---|---|---|
| MV E/A | 1.55±0.3 | 1.86±0.29** |
| MV E'/A' | 1.21±0.1 | 1.21±0.07 |
| MV E/E' | 28.4±4.9 | 26.92±4.8 |
| IVRT | 11.82±2.0 | 10.47±1.5 |
| SET (AET); ms | 50.29±3.06 | 47.99±3.75 |
| LV MPI | 0.44±0.06 | 0.39±0.051* |

**Table S1. Echocardiogram parameters from TgRRB and WT mice.** Systolic Measurements (N=6-8 mice per group) - HR: heart rate LVID; s: End-systolic Diameter of the left ventricle. LVID; d: End-diastolic Diameter of the left ventricle. LVAW; s: End-systolic Left Ventricular Anterior Wall thickness. LVPW; s: End-systolic Left Ventricular Posterior Wall thickness. LVAW; d: End-diastolic Left Ventricular Anterior Wall thickness. LVPW; d: End-diastolic Left Ventricular Posterior Wall thickness. EF: Ejection Fraction. FS: Fractional Shortening. LV MPI: Left Ventricle Myocardial Performance Index. Diastolic Measurements (N=13-14 mice per group) - MV E/A: Ratio between early(E) and late(A) trans mitral flow velocities. E/E': Mitral inflow E wave/tissue Doppler mitral annulus velocity ratio. IVRT: Isovolumic Relaxation Time. SET: systolic ejection time. P values were determined using unpaired, two-tailed, Student's t-tests. *p ≤ 0.05, **p ≤ 0.01 vs WT.

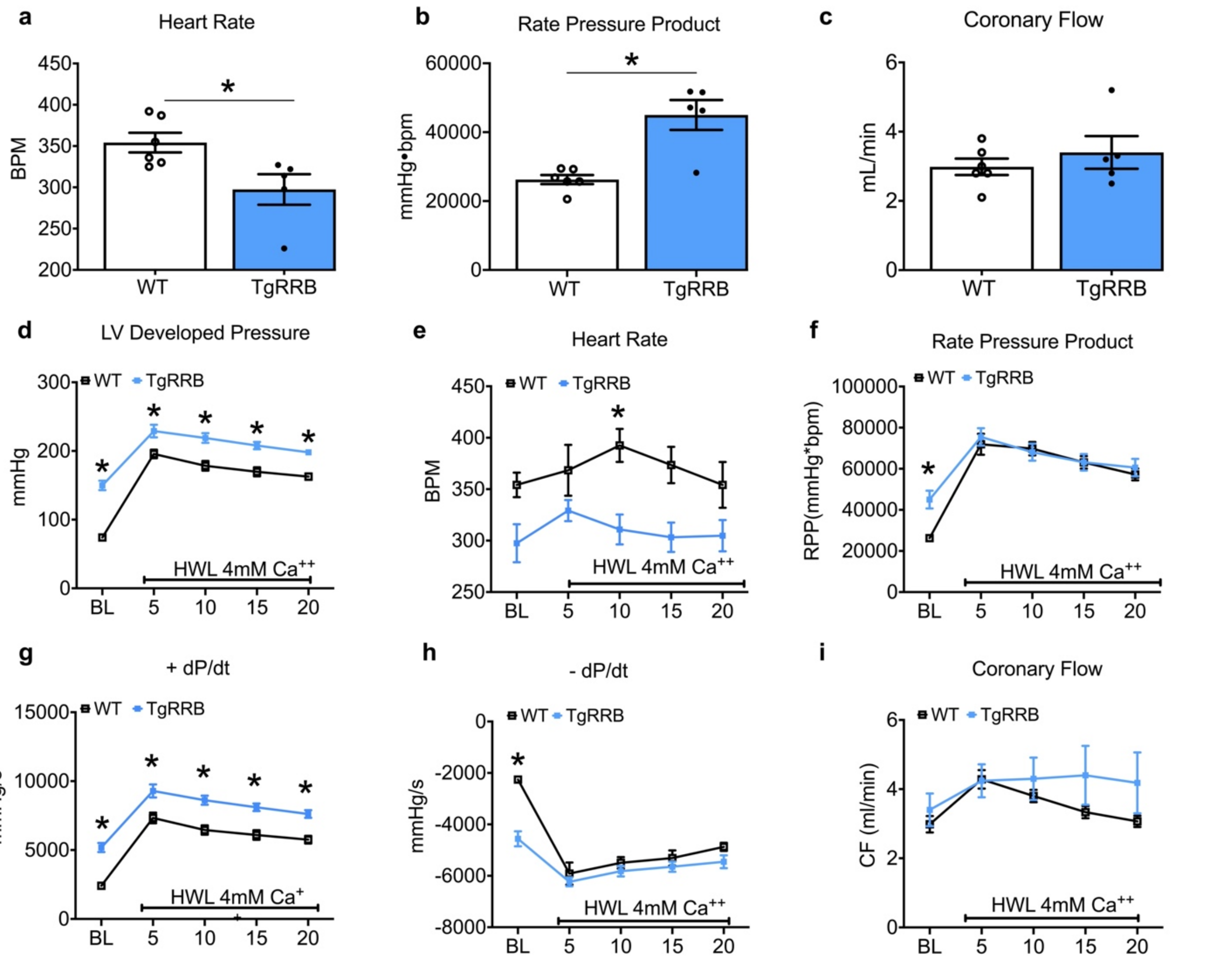


**Fig. S3: Contractility in Isolated perfused hearts at baseline and under high workload induced by elevating $Ca^{2+}$.** (a-c) Baseline measurements (a) spontaneous heart rate (b) Rate-pressure product (Cardiac output) (c) Coronary Flow. N = 6-7 mice per group. *$p \leq 0.05$ vs WT using unpaired t-test. (d-i) High workload measurements (d) Left ventricular developed pressure (e) Heart rate (f) rate-pressure product (g) +dP/dt (h) -dP/dt (i) Coronary N ≥ 10 mice per group. All data are presented as mean ± S.E.M. P values were determined using two-way ANOVA. *$p \leq 0.05$ vs WT.

**a**

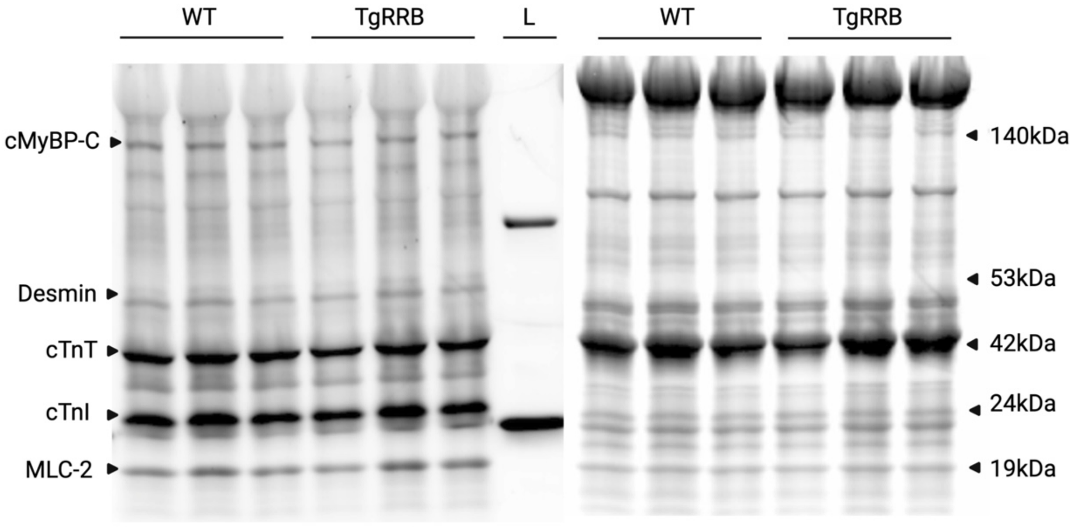


**b**

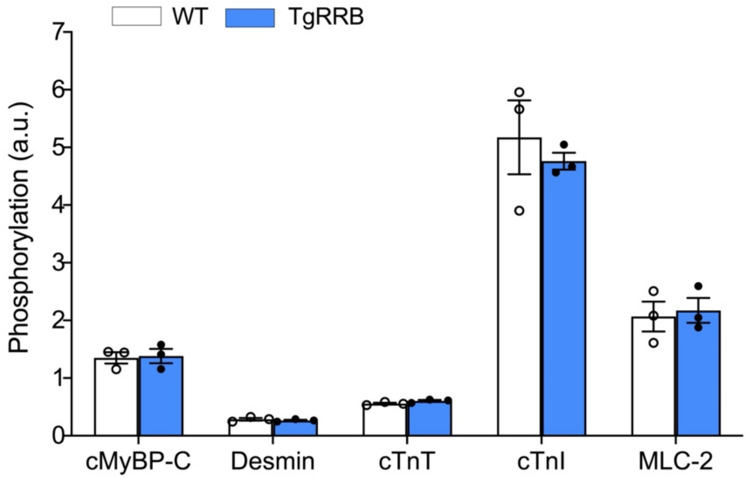


**Fig. S4. Chronic dATP elevation does not alter phosphorylation status of sarcomeric proteins.** (a) ProQ Diamond (left) and SYPRO Ruby (right) stained gels of isolated myofibril lysate of WT and TgRRB LV tissue with L (ladder) showing 25kDa and 75kDa bands from bottom (b) Densitometry analysis of the phosphorylation level normalized to protein levels. n=3 mice. All data are presented as mean ± S.E.M.

| Contractile function | WT-CMs | TgRRB-CMs |
|---|---|---|
| Fractional shortening (%) | 7.1±0.2 | 8.98±0.2* |
| Maximum rate of contraction (μm/s) | -3.6±0.1 | -4.5±0.1* |
| Time to peak (ms) | 83±0.1 | 77±0.4* |
| Maximum rate of relaxation (μm/s) | 2.4±0.1 | 3.5±0.1* |
| Time to 50% relaxation (ms) | 77±4 | 69±3 |
| Time to 90% relaxation (ms) | 142±7 | 119±6* |
| Calcium handling | WT-CMs | TgRRB-CMs |
| Resting $Ca^{2+}$ (Fura-2 ratio) | 1.86±0.01 | 1.87±0.01 |
| $Ca^{2+}$ transient amplitude (% of baseline Fura-2 ratio) | 27.9±1.4 | 31.6±1.66 |
| Maximum rate of $Ca^{2+}$ release (U/s) | 45.6±2.2 | 52±2 |
| Maximum rate of $Ca^{2+}$ decay (U/s) | -2.3±0.09 | -3.8±0.2* |
| Time to transient peak (ms) | 83±2.2 | 83±2 |
| Time to 50% transient decay (ms) | 77.1±4.1 | 69.2±3.4 |
| Time to 90% transient decay (ms) | 142±7 | 120±7* |
| Sarcoplasmic reticulum $Ca^{2+}$ load (F/F0) | 2.981±0.183 | 2.832±0.235 |

**Table S2. Contractile and Calcium transient properties of isolated CMs from TgRRB mice.** N = 6-7 mice per group, n≥70 cells per group. P values were determined using unpaired, two-tailed, Student's t-tests. *p ≤ 0.05 vs WT.

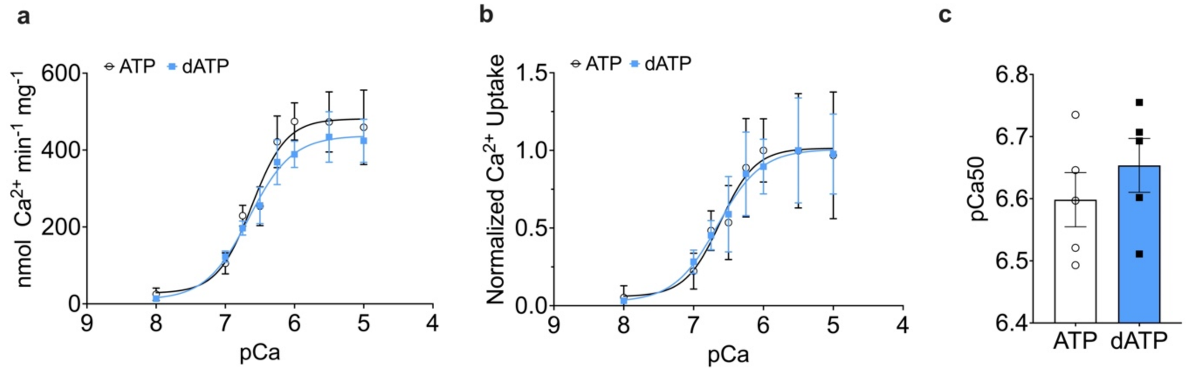


**Fig. S5. dATP does not alter $Ca^{2+}$ uptake in the sarcoplasmic reticulum.** (a) Uptake of $^{45}Ca^{2+}$ in mouse heart homogenate incubated in varying [$Ca^{2+}$] and 5 mM ATP (black) or dATP (blue). (b) Normalized $Ca^{2+}$ uptake and (c) $Ca^{2+}$ concentration at half-maximal activity (pCa50) are not changed with dATP. n=5 mice. All data are presented as mean ± S.E.M.

Given that the maximum rate of calcium decay is increased in both transgenic and transduced CMs, we hypothesized that maximum $Ca^{2+}$ uptake by SERCA might be increased with elevated dATP. This idea is supported by previously published findings by Trumble, Sutko, and Reeves that dATP can be used as a substrate by SERCA2a in isolated sarcoplasmic reticulum vesicles and increases maximal pump activity[1]. To verify this hypothesis in cardiac tissue, C57/BL6 mouse hearts were homogenized and incubated with $^{45}Ca$ in buffers containing either ATP or dATP in buffers with pCa varying from 8 to 5 and radioactivity in the homogenate was measured using a scintillation counter. While results confirm that SERCA2 can use dATP as a substrate with similar affinity to ATP, neither total uptake (473±78 for ATP, 434±65 for dATP, arbitrary units normalized to total protein) or calcium sensitivity (pCa50, 6.59±0.03 with ATP, 6.70±0.16 with dATP) were significantly changed with nucleotide.

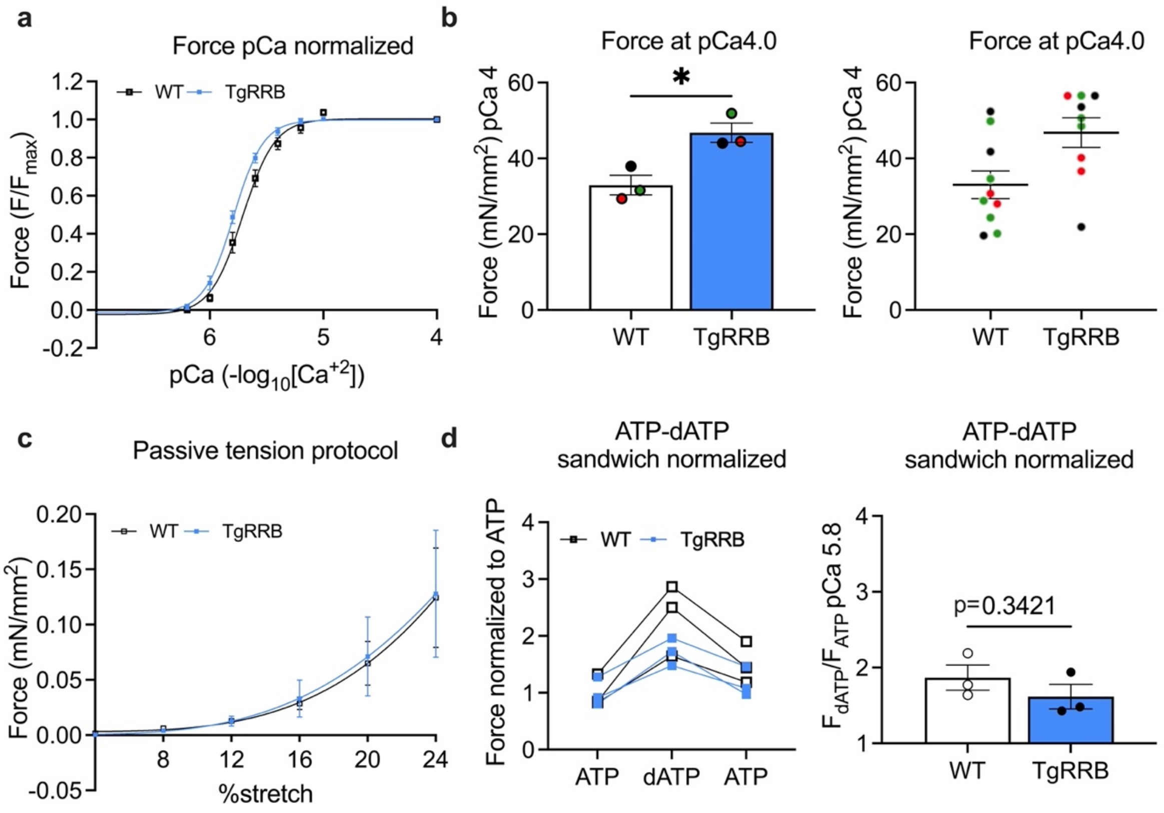


**Fig. S6. TgRRB trabeculae demonstrated similar passive stiffness and greater force generation than WT independent of dATP. (a) Normalized** force vs. pCa of demembranated trabeculae from WT and TgRRB hearts (b) Average values of Force measured at pCa4 per animal (right) and technical replicates of Force measured at pCa4 (left) (N=3 animals/group). (c) Passive force in resting conditions in response to stretch of trabeculae from WT and TgRRB hearts (d) Normalized force generated by skinned trabeculae from WT and TgRRB hearts in presence of ATP vs. dATP (left) and the bar graph depicting average values (N=3 animals/group). All data are presented as mean ± S.E.M. P values were determined using unpaired, two-tailed, Student's t-tests. *P<0.05.

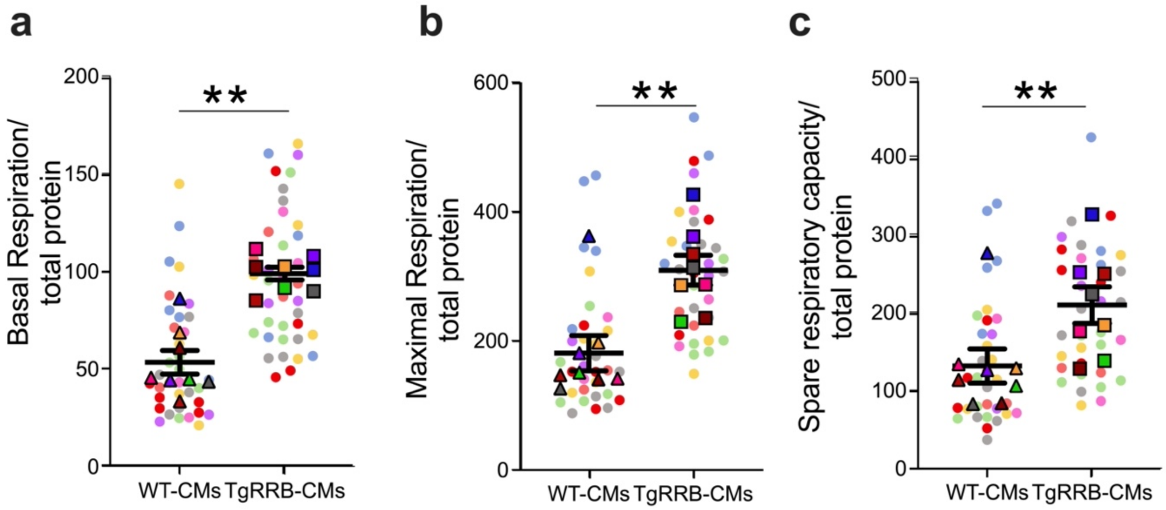


**Fig. S7. Individual data points for the Oxygen consumption rate (OCR) measurements in TgRRB-CMs vs WT-CMs.** OCR was measured on a standard Seahorse Mito stress test. (a) Basal OCR (b) Maximal OCR and (c) Spare respiratory capacity normalized to the total protein. The solid symbols are averaged values of technical replicates (lighter dots) per animals (N=8 animals/group). All data are presented as mean ± S.E.M. P values were determined using unpaired, two-tailed, Student's t-tests. *P<0.05, **P<0.01.

**a**

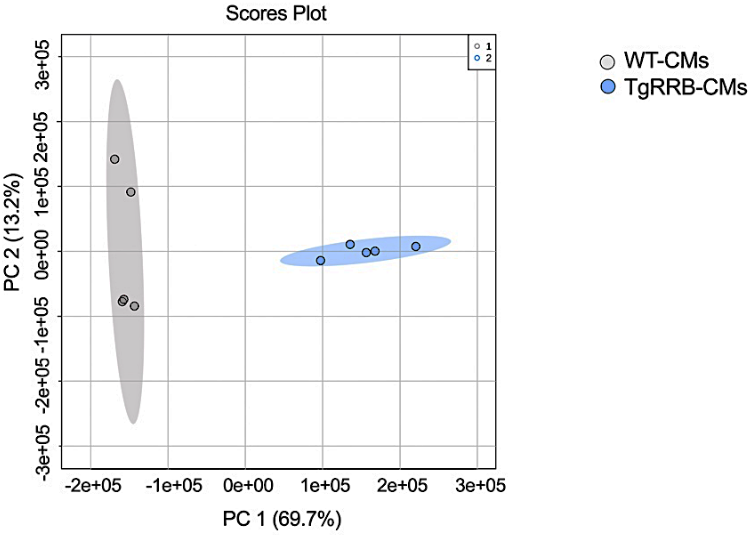


**b**

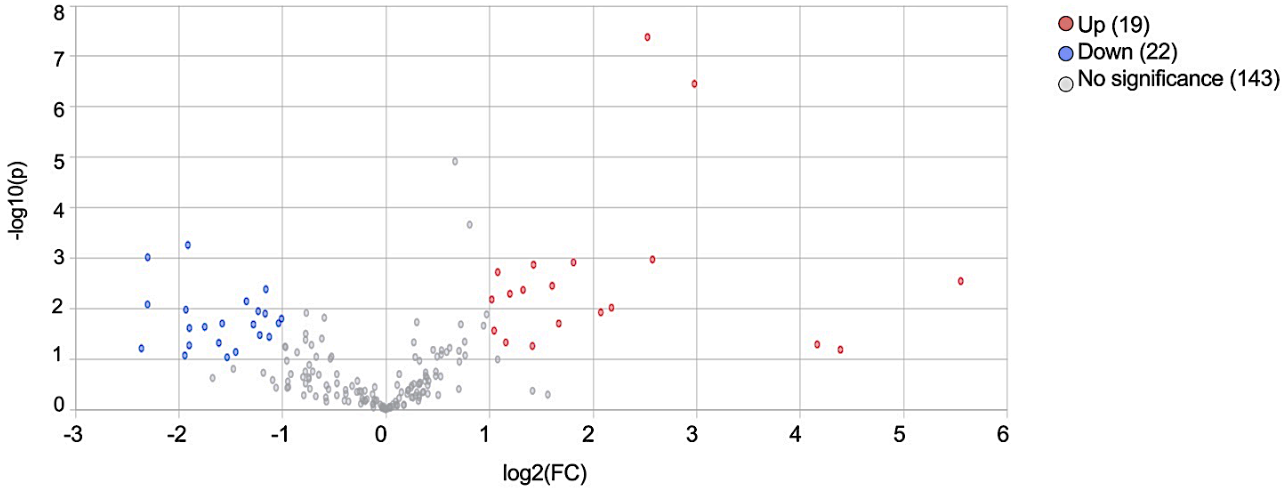


**Fig. S8 Metabolomic profile of TgRRB-CMs.** Metabolomic analysis was performed on isolated cardiomyocytes from TgRRB mice and age-matched control mice using LC/MS. (a) PCA of all 184 metabolites clearly separates the profiles of TgRRB-CMs (blue) and WT-CMs (grey).
(b) Volcano plot of the log of fold change in metabolites between TgRRB-CMs and WT-CMs group. Statistical analysis was performed by two-tailed F-test for differential metabolite expression analysis showed 19 upregulated and 22 downregulated metabolites. (n=5 for TgRRB mice and n=5 for control WT mice)

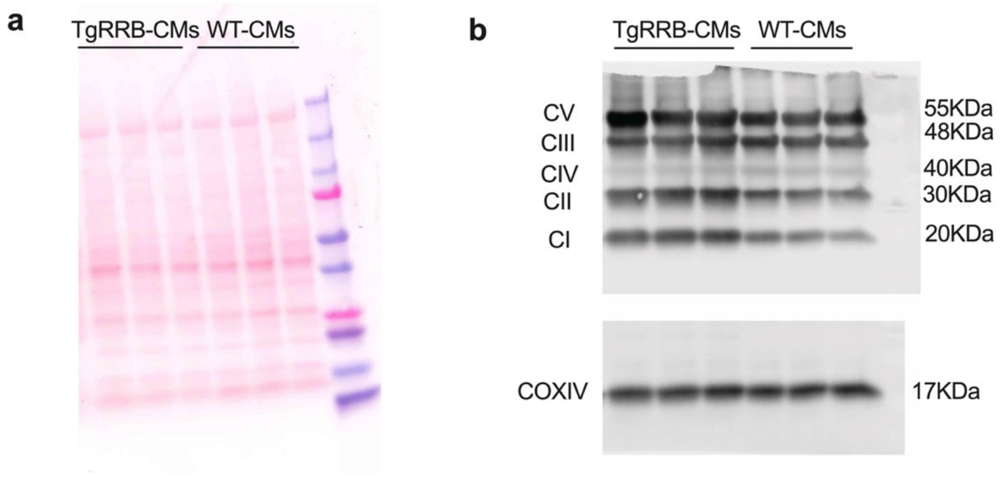


**Fig. S9. OXPHOS Western blot.** (a) Ponceau staining (b) OXPHOS (ETC complex) immunoblot of proteins CV (ATP5A), CIII (UQCRC2), CIV (MTCO1), CII (SDHB) and CI (NDUFB8) with COXIV protein as a loading control. N=3 animals per group.

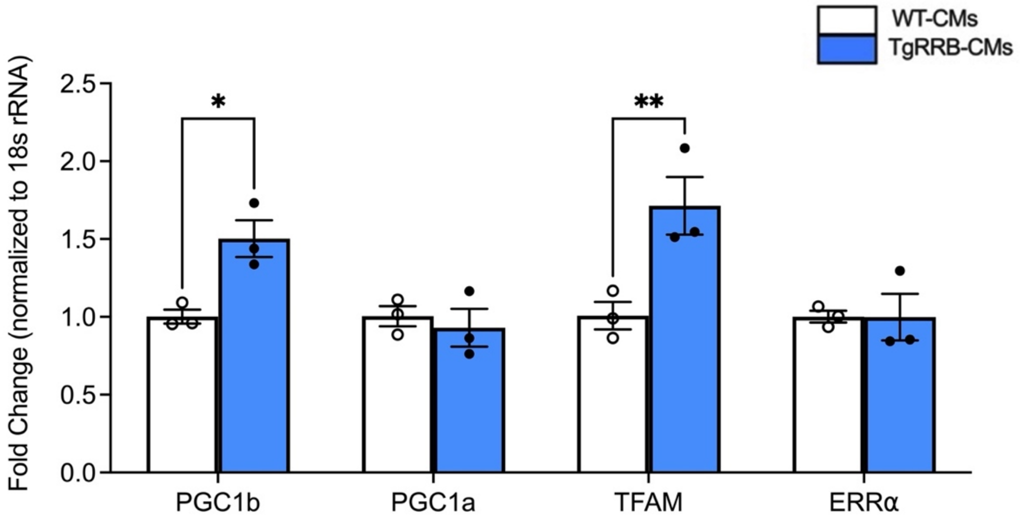


**Fig. S10. TgRRB hearts demonstrated greater expression of genes involved in mitochondrial biogenesis and turnover.** qRT-PCR analyses of of indicated genes involved in mitochondria biogenesis = (N=3 animals/group). All data are presented as mean ± S.E.M. P values were determined using unpaired, two-tailed, Student's t-tests. *P<0.05, **P<0.01.

### Methods

#### Myofilament protein phosphorylation

Protein phosphorylation was determined as previously described[2]. Isolated myofibril lysates were separated on gradient gels (Criterion Tris– HCl 4–20% gel, BioRad) and phosphorylated proteins were detected with Pro-Q Diamond staining (Life Technologies, Darmstadt, Germany). For protein content, gels were stained with SYPRO-Ruby (Life Tech- nologies, Darmstadt, Germany). Phosphorylation of myofilament proteins was determined relative to protein expression.

#### SR calcium uptake assay

Ventricles from adult mice were homogenized in ice-cold buffer (mM: 50 $KH_2PO_4$, 10 NaF, 1 EDTA, 300 sucrose, 0.3 PMSF, 0.5 DTT). Homogenate was added to uptake buffer (mM: 0.5 EGTA, 0.5 MgCl2, 5 MgATP, 10 creatine phosphate, 40 imidazole, 5 potassium oxalate, 5 NaN3, 10 procaine, 0.03 ruthenium red, pH 7.1) containing calcium at concentrations varying from pCa 8 to 5. After a two-minute pre-incubation of the assay buffer containing 150 µCi/mL $^{45}Ca$ (Perkin Elmer, Waltham, MA) at 37°C, the ventricular homogenate was incubated with continuous stirring. The reaction was stopped after two minutes by filtration through a 0.45-µM Millipore filter and washed twice with cold buffer (20 mM Tris, 2mM EGTA pH 7.0). Radioactivity was measured using a scintillation counter and normalized to protein content in the homogenate determined by Pierce BCA protein assay. Total vesicular calcium uptake was calculated by determining the amount of $^{45}Ca$ bound to the Millipore filters via a liquid scintillation counter. The following equation was used to determine the SR $Ca^{2+}$ uptake rate: $Ca^{2+}$ Uptake = nmol Ca x (counts per minute/ (specific activity x 5) x 1/mg protein in 300 µL) x 0.5.

Permeabilized trabeculae mechanics

Experiments were performed as described in [3] and [4]. In brief, excised hearts were demembranated overnight at 4°C in a relaxing solution (in mM: 100 KCl, 10 imidazole, 2 EGTA, 5 MgCl2, and 4 ATP) containing 50% glycerol (vol:vol), 1× protease inhibitor cocktail (Sigma-Aldrich, P8340), PhosSTOP phosphatase inhibitor cocktails (Sigma-Aldrich), and 1% Triton X-100. Permeabilized trabeculae were then dissected from dissected out and wrapped between aluminum t-clips, then mounted between a motor (Aurora Scientific, Model 312B) and force transducer (Aurora Scientific, Model 403A) using custom aluminum T-clips. Preparations were moved to a bath containing experimental relaxing solution at pH 7.0 and 15°C containing (in mM): 15 phosphocreatine, 15 EGTA, 80 MOPS, 1 free $Mg^{2+}$, $10^{-6}$ $Ca^{2+}$, 1 DTT, and 5 $Mg^{2+}$ ATP. SL was measured using a Fourier transform of a digitized image of the sarcomeres using an IonOptix camera connected to a ×40 dry objective lens. SL was set to 2.3 μm for the experiments. Trabeculae were submerged in physiological solution at 15°C containing a range of pCa (= –log [$Ca^{2+}$]) from 9.0 to 4.0 and allowed to reach steady state at each pCa. Force-pCa curves for each genotype were collected and analyzed using custom code with LabView software.

For passive tension measurements, preparations were set to length just above slack (L0), then sequentially stretched at 4% increments up to 24% L0. We calculated median passive force (mN/mm2) per mouse from multiple preparations (1–4 per heart). We scaled all data to controls at 24% length change for each experiment to compare 3 experiments at once. N = 3 mice per group.

Quantitative Real-Time PCR

Total DNA preparations were obtained by using the DNAeasy blood and tissue kit protocol (Qiagen). Quantitative real-time PCR, for mtDNA copy number, expression of mitochondria encoded NADH dehydrogenase 1 gene and endogenous control 18S rRNA gene were measured on an ABI PRISM 7900HT Sequence Detection System (Applied Biosystems). The forward and reverse primers for PCR amplification are been listed in[5].
